# Comparative functional genomics of six *Pectobacterium* species reveals core and unique fitness determinants in potato tubers

**DOI:** 10.64898/2026.10.02.755747

**Authors:** Tyler C. Helmann, Mary Ann Karp, Margery Daughtrey, Lindsey E. Reid, Paul V. Stodghill, Melanie J. Filiatrault

**Author notes:** Corresponding author: M. J. Filiatrault, and. Funding: U.S.D.A Agricultural Research Service: 8062-21000-048-000-D.

## Abstract

*Pectobacterium*, a member of the soft rot Pectobacteriaceae (SRP), can cause highly destructive soft rot and blackleg diseases on many important crop and other host plants, particularly potato (*Solanum tuberosum*). *Pectobacterium* and its sister genus *Dickeya* have common virulence strategies, yet a highly diverse “open” pangenome containing many unique and hypothetical proteins. Here, we used randomly-barcoded transposon insertion-site sequencing (RB-TnSeq) on six *Pectobacterium* species to identify genes involved in bacterial survival and growth in potato tubers, as well as in minimal media conditions containing different single carbon sources. The strains evaluated varied broadly in genomic content as well as host plant and year of isolation. We focused on identifying conditionally essential genes across all strains tested, as well as genes that were important for growth in one or few of the tested strains. These results highlight the importance of many common genes involved in metabolism and maintaining cellular homeostasis, as well as shared and rare virulence traits including polygalacturonate catabolism, genes involved in polysaccharide biosynthesis, immunity proteins, and (for *P. brasiliense*) self-tolerance of the antibiotic carbapenem. This large dataset of conditionally essential genes, generated across *Pectobacterium* species during soft rot infection of the agriculturally important plant host potato, can serve as a rich dataset for further hypothesis generation and identification of important and/or novel *in planta* fitness factors.

## Introduction

Bacteria in the genera *Pectobacterium* and *Dickeya* are damaging soft rot pathogens of potato (*Solanum tuberosum*) and a broad range of other crop plants. These necrotrophic pathogens utilize a repertoire of secreted plant cell wall degrading enzymes (PCWDEs) to release host nutrients and access complex carbon-containing polymers such as pectins (Toth et al., 2003; Charkowski et al., 2012). Though these genera are highly diverse, with the *Pectobacterium* genus having an open pangenome (Zoledowska et al., 2018; Jonkheer et al., 2021) and comprising at least 20 species (Sawada et al., 2024), important traits for host colonization have been described and include commonly cited virulence factors such as the quorum-sensing coordinated production and secretion of high levels of PCWDEs (Toth et al., 2003; Liu et al., 2008). Despite this shared functionality within the “core” genome, there are thousands of genes that are maintained in a species- or strain-specific manner with unknown contributions to bacterial fitness in the environment.

Because most *Pectobacterium* species are pathogenic on potato, available genome sequences are disproportionately derived from potato-associated isolates that tend to be pathogenic (Toth et al., 2021). This sampling bias limits comparative genomic analyses of virulence-related and environmental survival traits across the genus. There are a wide variety of genes present in the broader pangenomic gene pool, and much less is known about the non-disease state lifestyles of these organisms, which can also survive in the soil and groundwater, as well as on plants in a latent or dormant phase (Toth et al., 2021). Plant pathogens like *Pectobacterium* spp. are commonly found in diverse environments and must compete for nutritional resources with other microorganisms during the infection of host plants. In fact, field infections are often observed in which multiple soft rot pathogens are present, and it is unclear if these interactions are more competitive or synergistic (Motyka-Pomagruk et al., 2020; Ge et al., 2021).

Previous studies have used comparative genomics to identify orthologous genes for the genus *Pectobacterium*, including functional prediction and identification of variation between species (Glasner et al., 2008; Li et al., 2018; Zoledowska et al., 2018; Arizala and Arif, 2019; Bellieny-Rabelo et al., 2019; Jonkheer et al., 2021). However, there are challenges for identifying protein function based on homology-based predictions, particularly in cases where genes are not widely present in related strains. In this study, we leveraged high-throughput functional genomic screening to test for essential and conditionally essential genes (i.e. dispensable in optimal growth conditions and essential or having a growth defect in other growth conditions) across multiple species during infection of the agriculturally-important plant host potato. By measuring relative abundance of random insertional mutants before and after population growth in a given experimental condition, high-throughput functional screening methods like transposon insertion-site sequencing (TnSeq) are valuable tools to identify genes that are essential or conditionally essential under specific growth conditions (Cole et al., 2017; Helmann et al., 2020; Georgoulis et al., 2021; Luneau et al., 2022). Our recent studies using randomly-barcoded TnSeq (RB-TnSeq) in *Dickeya dadantii* and *D. dianthicola* generated genome-wide mutant fitness profiles showing that successful growth in potato depends on both core metabolic functions and more specialized traits, with genes involved in amino acid biosynthesis, stress responses, iron acquisition, motility, chemotaxis, and plant cell wall degradation making major contributions to fitness in tubers and stems (Helmann et al., 2022; Gonzalez-Tobon et al., 2024). Other TnSeq studies in *D. dadantii* and *D. solani* have identified conserved genes broadly important for plant host colonization and virulence, with some variation by host tissue and the environment (Royet et al., 2019; Robic et al., 2023). Several studies have generated and characterized individual transposon mutants in *Pectobacterium*, identifying important genes required for potato virulence primarily basic metabolism, extracellular enzyme production (i.e. PCWDEs), and biofilm formation (Hinton et al., 1989; Lee et al., 2013). In this study, we constructed and used RB-TnSeq in six *Pectobacterium* species to identify conditionally essential or important genes for growth in potato tubers, as well as both rich and minimal media. We screened well-studied model *Pectobacterium* strains alongside some other plant-associated *Pectobacterium* isolates, including those from potato, petunia, and calla lily. Although these strains are expected to share core metabolic requirements and virulence strategies, we sought to identify strain-specific conditionally essential genes. Using multiple barcoded transposon mutant libraries across *Pectobacterium* spp. we generated a broad dataset of conditionally essential genes across strains and ecologically-relevant conditions, including growth in potato tubers.

## Results

### Genomic characteristics of *Pectobacterium* strains used in this study

*Pectobacterium* spp. are necrotrophic pathogens capable of infecting a diverse range of plant hosts, including both dicotyledonous and monocotyledonous species. However, a major challenge in leveraging their genomic diversity is that the vast majority of sequenced strains have been isolated from potato, an economically important host. To address this sampling bias and better capture the diversity of *Pectobacterium* associated with alternative hosts, we isolated two additional strains from non-potato host plants. *Pectobacterium aroidearum* 61-19 (Par6119) was isolated from calla lily (*Zantedeschia*) and *Pectobacterium brasiliense* 123-1 (Pbr123) was isolated from petunia (*Petunia*). Following identification to species, we performed whole-genome sequencing, assembly, and annotation. We also re-sequenced and closed the genomes of two isolates from NY potato previously identified and published in (Ma et al., 2021): *Pectobacterium parmentieri* NY1722A (PpaNY1722A) and *Pectobacterium versatile* NY1715C (PveNY1715C). Whole-genome sequencing and annotation metrics are listed in Table 1. The plasmid identified in PpaNY1722A measures 8,291 bp and contains ten annotated genes (AC4LGQ_23585 - AC4LGQ_23630). We performed analyses of these four strains alongside the well-characterized model strains *Pectobacterium atrosepticum* SCRI1043 (PatSCRI1043), *Pectobacterium brasiliense* 1692 (Pbr1692), and *Pectobacterium carotovorum* WPP14 (PccWPP14).

**Table 1.**
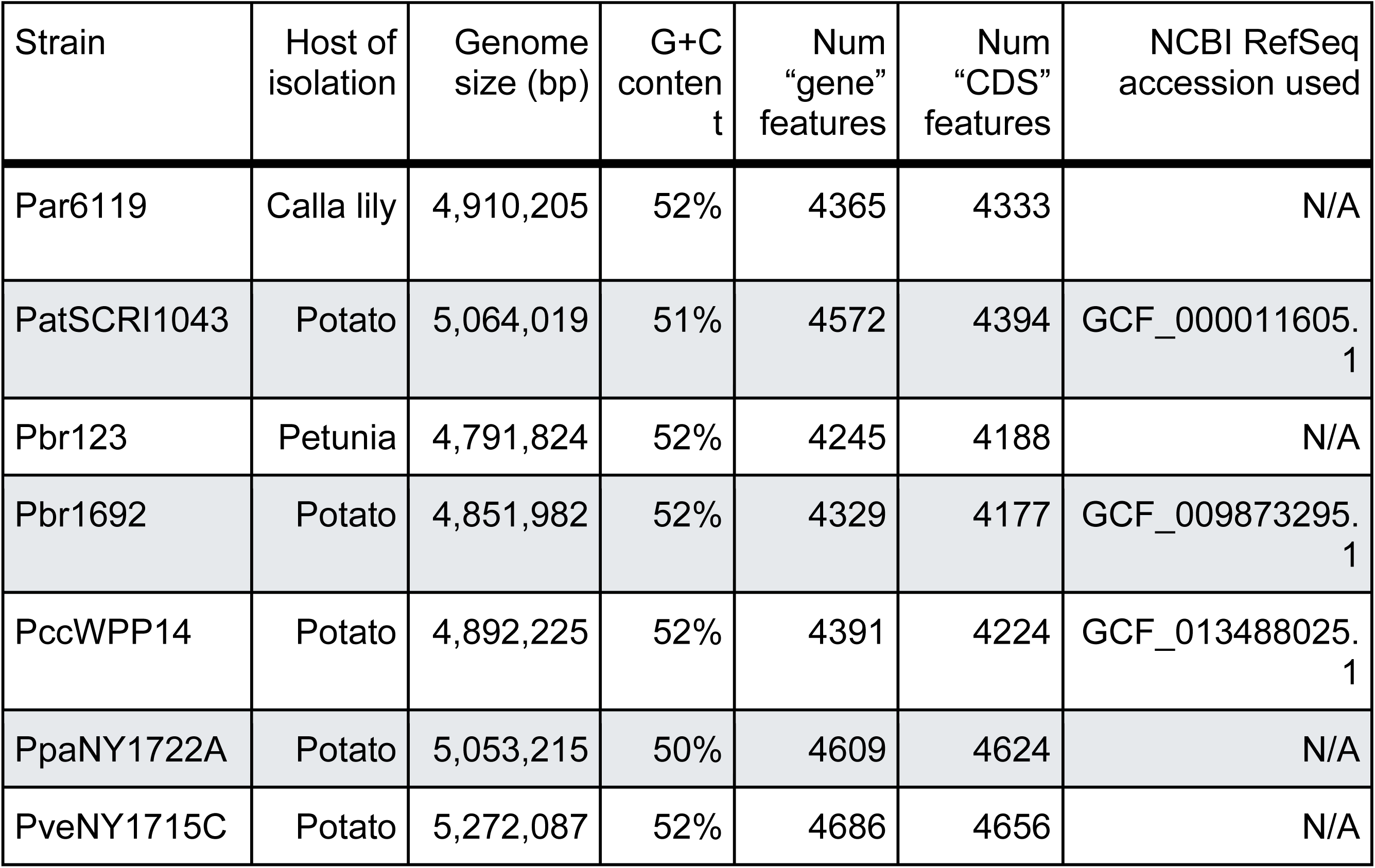
Genome statistics of the seven *Pectobacterium* strains used this study. The genomes were annotated by NCBI using PGAP (Tatusova et al., 2016). Strain PpaNY1722A is unique among these strains in that it contains a plasmid measuring 8,291 bp.

### RB-Tn library construction

We constructed barcoded transposon insertional mutant libraries in seven *Pectobacterium* strains comprising six species. We focused on generating mutant libraries in a mix of model organisms (PatSCRI1043, Pbr1692, and PccWPP14) and recent environmental isolates from both potato (PpaNY1722A and PveNY1715C) and other plants (Par6119 and Pbr123). Due to strain-specific variation in transformation efficiency, we tested both Tn5 and *mariner* barcoded transposon donor plasmids from (Wetmore et al., 2015) for the target recipient strains. For Par6119 and PpaNY1722A Tn5 resulted in more insertions, while *mariner* resulted in more insertions for PatSCRI1043, Pbr123, Pbr1692, PccWPP14, and PveNY1715C (data not shown). The resulting mutant libraries ranged in size from ∼185k to ∼582k strains, and all libraries had central insertions (defined as the middle 10-90% of the coding region) in >90.7% of total annotated genes in the genome (Table 2). Insertions within all chromosomes and the PpaNY1722A plasmid were evenly distributed, as evidenced by the overall gene coverage as well as calculated insertion density (Supplementary Figure 1). For all strains, we used eggNOG-mapper v2.1.12 (Cantalapiedra et al., 2021) for further functional annotation to assign protein sequences to COG (Clusters of Orthologous Groups) categories. Using the per-gene transposon insertion densities, we identified 315 to 399 genes per strain that are likely to be essential for growth on LB (or simply, “essential”) based on a lack of central insertions (Supplementary Table 2) during library construction.

**Table 2.** Insertion statistics for *Pectobacterium* randomly-barcoded transposon libraries.

| Library | Transposon | Insertions in genome | Protein-coding genes with central insertions (Total) | Median strains per hit protein |
| --- | --- | --- | --- | --- |
| Par6119 | Tn5 | 359,157 | 3931 (4293) | 26 |
| PatSCRI1043 | Mariner | 240,722 | 4023 (4392) | 27 |
| Pbr123 | Mariner | 388,212 | 3832 (4156) | 45 |
| Pbr1692 | Mariner | 185,037 | 3789 (4174) | 21 |
| PccWPP14 | Mariner | 284,613 | 3860 (4220) | 34 |
| PpaNY1722A | Tn5 | 586,547 | 4188 (4574) | 39 |
| PveNY1715C | Mariner | 511,292 | 4252 (4574) | 52 |

### Ortholog analysis of the *Pectobacterium* genus

To allow for direct comparisons of gene fitness data between strains, OrthoFinder v3 (Emms et al., 2026) was used to identify orthologous genes or orthogroups (OGs) among “Complete” and “Chromosome” level *Pectobacterium* assemblies from NCBI RefSeq (O’Leary et al., 2016) (N=151), and the four sequenced strains described above: Par6119, Pbr123, PveNY1715C, and PpaNY1722A. We used this expanded dataset to better provide context for OrthoFinder clustering of the proteomes of these strains. Of 655,344 total genes analyzed, 651,999 (99.5%) were assigned to 9,539 orthogroups, including 1,114 single-copy orthogroups (Supplementary Table 1). The distribution of the 9,539 identified orthogroups was as follows: 2,763 “core” orthogroups present in >95% (>147) total strains; 2,836 “shell” orthogroups present in ≥15% and ≤95% (15-147) total strains; and 3,940 “cloud” orthogroups present in <10% (<15) total strains (Figure 1).

**Figure 1.**
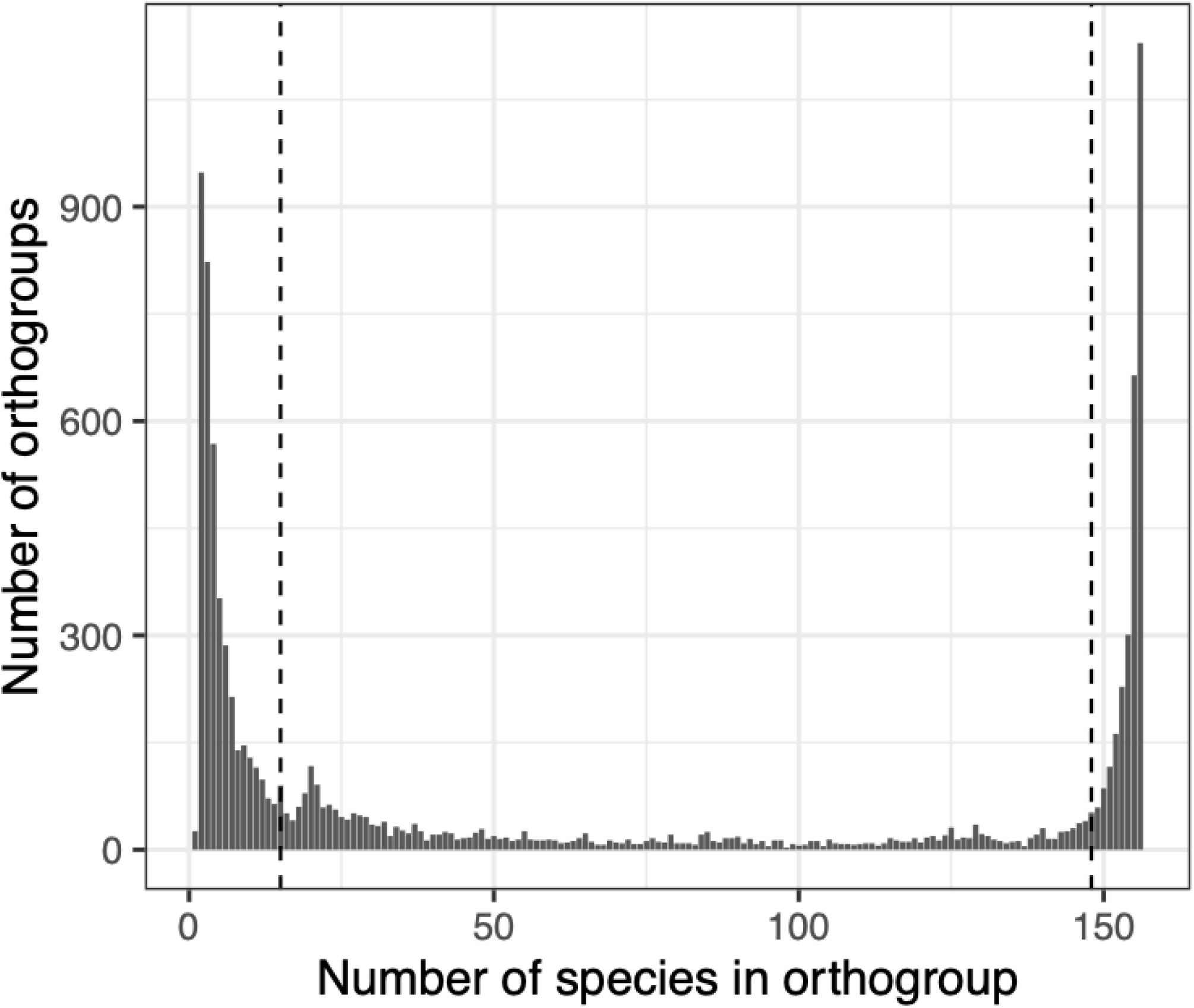
*Pectobacterium* pangenome U-plot of orthogroup distributions. OrthoFinder v3 (Emms and Kelly, 2019) was run on 155 *Pectobacterium* proteomes representing 21 different species. Dashed vertical lines are shown at cloud/shell cutoff = 15 (10%) and shell/core cutoff = 147 (95%). Plot was generated using “ggplot2” (Wickham, 2016) in R.

### Orthogroup identification facilitates analysis of core genome and shared essential genes

Based on the calculated orthogroups (Supplementary Table 1), we identified 3,000 orthogroups containing genes present in all seven strains, including 228 shared likely essential genes (Supplementary Figure 2). As expected, some of the genes consistently predicted to be essential are those predicted to encode 30S ribosomal proteins (N=19), 50S ribosomal proteins (N=24), cell division proteins FtsAILQWYZ, DNA polymerase III subunits (N=5), RNA polymerase subunits (alpha, beta, beta’), sigma factors RpoD and RpoH, and tRNA ligases (N=22). Even between Pbr1692 and Pbr123, member strains of the same species, there was substantial variation in shared genes. For example, while Pbr1692 and Pbr123 share 3,571 orthogroups, Pbr1692 has 422 and Pbr123 has 441 genes that are not found in the other strain. Of those, 112 (27%) and 115 (26%) (Pbr1692 and Pbr123, respectively) are annotated as encoding hypothetical proteins. More broadly, 173 orthogroups are found only in Pbr1692 and not any of the other six strains examined here, and 73 (42%) of those genes are annotated as encoding hypothetical proteins.

### Overall results of BarSeq fitness assays

For each transposon library constructed, we tested relative mutant fitness by growing the pooled populations in rich media (LB), minimal media (M9) containing various carbon sources, and stab-inoculated potato tubers. Four strains were also tested in LB under heat stress at 36°C; specifically Pbr123, Pbr1692, PccWPP14, and PveNY1715C. *P. atrosepticum* and *P. parmentieri* can only grow at temperatures up to 33°C (Bastas, 2023). We used barcode sequencing (BarSeq) to calculate per-gene fitness values (following standard analyses (Wetmore et al., 2015)) for 3,557 to 3,944 genes per strain, representing 82 to 86% of the genome for each strain. Overall, between 31 and 61 experimental samples (not including Time0 samples) per strain passed previously established QC metrics (Wetmore et al., 2015) and were uploaded to the Fitness Browser (https://fit.genomics.lbl.gov/). Supplementary Table 3 summarizes the breakdown of replicates by strain and condition. Although experiments performed in the heat shock condition were sequenced and processed as part of this dataset, discussion of this treatment will be described separately in future work.

In each experimental sample, per-gene fitness was calculated as a weighted average of potentially many insertional mutant strains within each gene. For initial analysis, we examined the lists of all genes that passed previously-established cutoffs for a strong phenotype; that is, an absolute fitness value >2 and absolute t-like test statistic >5 (Wetmore et al., 2015). Fitness is calculated on a log2 scale, and for example a fitness value of −2 indicates the mutant grew ∼25% as well as wild-type. These cutoffs were applied to all experimental samples independently, and results are shown in Supplementary Table 4. We focused our analyses on genes with significant phenotypes in multiple replicates of a given experimental condition and (in most cases) orthogroups having significant phenotypes across multiple strains. With the aim of identifying important functional gene categories, we used eggNOG-mapper (Cantalapiedra et al., 2021) to assign COG functional categories to all genes in each strain.

To further reduce gene fitness noise due to population stochasticity, especially *in planta*, we calculated the mean fitness value for each gene across replicate conditions tested (N= 2-3 *in vitro* and 4-9 *in planta*) (Supplementary Table 5). We confirmed that replicate samples for each condition clustered using principle component analysis (Supplementary Figure 3). We then used these mean fitness values to identify genes with consistent and moderate to strong phenotypes [absolute(mean fitness) >1] for each strain.

We combined per-condition mean gene fitness values with orthogroup information to identify genes (as orthogroups) that were commonly conditionally essential during growth in minimal media, during growth and infection of potato tubers, and those genes that were species specific or otherwise more variable in contribution to competitive fitness. Furthermore, we also specifically compared the results for the libraries constructed in Pbr1692 and Pbr123; while these strains are the same species, we reasoned that having different original hosts of isolation could represent divergent virulence or environmental growth strategies.

### Core metabolic genes important for growth in minimal media

To generate a baseline dataset of genes important for growth and catabolism of common sugars and other carbon sources, we compared results from testing population growth in rich media (LB) and minimal media (M9) containing various carbon sources. Initial results, based on individual experimental samples and summarized by COG assignment, highlighted genes within the categories “amino acid transport and metabolism”, “nucleotide transport and metabolism”, and “coenzyme transport and metabolism” as contributing significantly to growth in minimal media (Figure 2, Supplemental Table 4).

**Figure 2.**
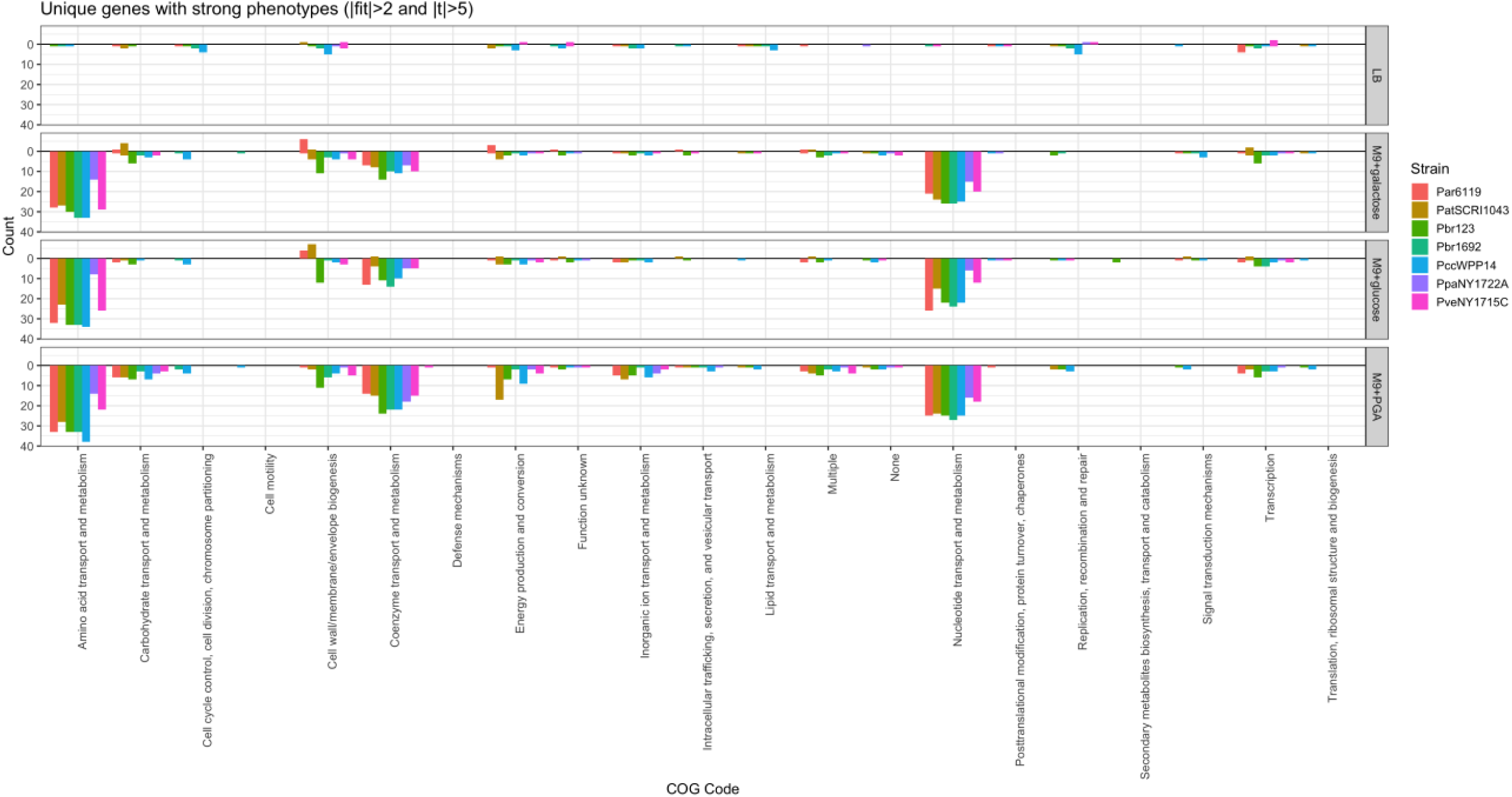
Number of unique genes with “strong” (|fitness|>2 and |t|>5) phenotypes in LB and M9 supplemented with galactose, glucose, or polygalacturonate. Direction of bar indicates a positive (up) or negative (down) fitness value. Plot was generated using “ggplot2” (Wickham, 2016) in R.

In the minimal media conditions, conditionally essential genes generally fell into two functional categories: 1) genes involved in amino acid, nucleotide, or coenzyme biosynthesis that were important in all minimal media experiments regardless of carbon source; and 2) catabolic genes that were specific to the carbon source being tested, including degradation products. For example, *argG* (HER17_RS21055^1^) encoding argininosuccinate synthase and *argH* (HER17_RS20620) encoding argininosuccinate lyase were highly important in all minimal media conditions tested, in all strains. In contrast, the genes encoding xylose isomerase (*xylA*, HER17_RS21085) and xylulokinase (*xylB*, HER17_RS21090) were required only for the utilization of xylose as a carbon source. The galactokinase-encoding gene *galK* (HER17_RS06390) was highly important for competitive fitness in both polygalacturonate and galactose conditions (Figure 3).

**Figure 3.**
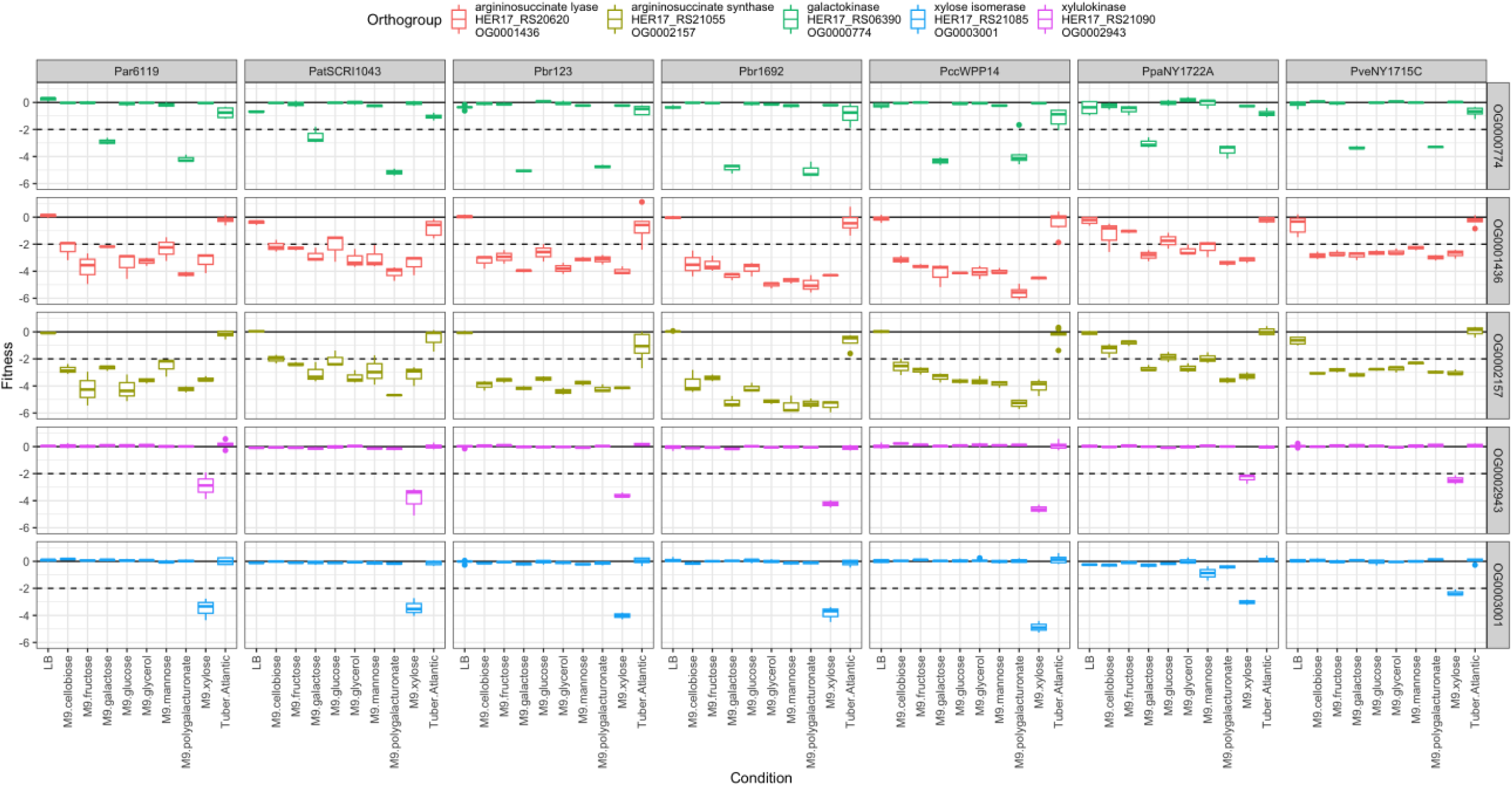
Fitness values across all strains tested for *argG* (HER17_RS21055), *argH* (HER17_RS20620), *galK* (HER17_RS06390), *xylA* (HER17_RS21085), and *xylB* (HER17_RS21090). The dashed line at fit = −2, is the default cutoff for a “strong” growth defect phenotype in the FEBA analysis pipeline, and indicates the mutant grew ∼25% as well as wild-type. Plot was generated using “ggplot2” (Wickham, 2016) in R.

Using an absolute(mean fitness) cutoff >1, we identified 35 genes important for growth in all seven strains tested in M9 with glucose (Figure 4), primarily amino acid transport and metabolism (N=19), nucleotide transport and metabolism (N=9), and coenzyme transport and metabolism (N=5). For growth in M9 with polygalacturonate, we identified 72 genes important in all 7 strains (Figure 4): amino acid transport and metabolism (N=25), nucleotide transport and metabolism (N=22), coenzyme transport and metabolism (N=16), carbohydrate transport and metabolism (N=2), and transcription (N=2). Of the 72 common genes important for growth in M9 with polygalacturonate, 35 of those were also important in M9 with glucose, highlighting the general requirements for growth in minimal media, and the additional complexity of catabolism of polygalacturonate relative to the simple sugar glucose. For example, common genes specific to catabolism of polygalacturonate included genes specific to polygalacturonate such as *kdgK* (HER17_RS00410), *kduD* (HER17_RS11410), and *uxaC* (HER17_RS18715), as well as related pathway genes more broadly important in the catabolism of galactose, like *galK* and *galT* (HER17_RS06390, HER17_RS06385).

**Figure 4.**
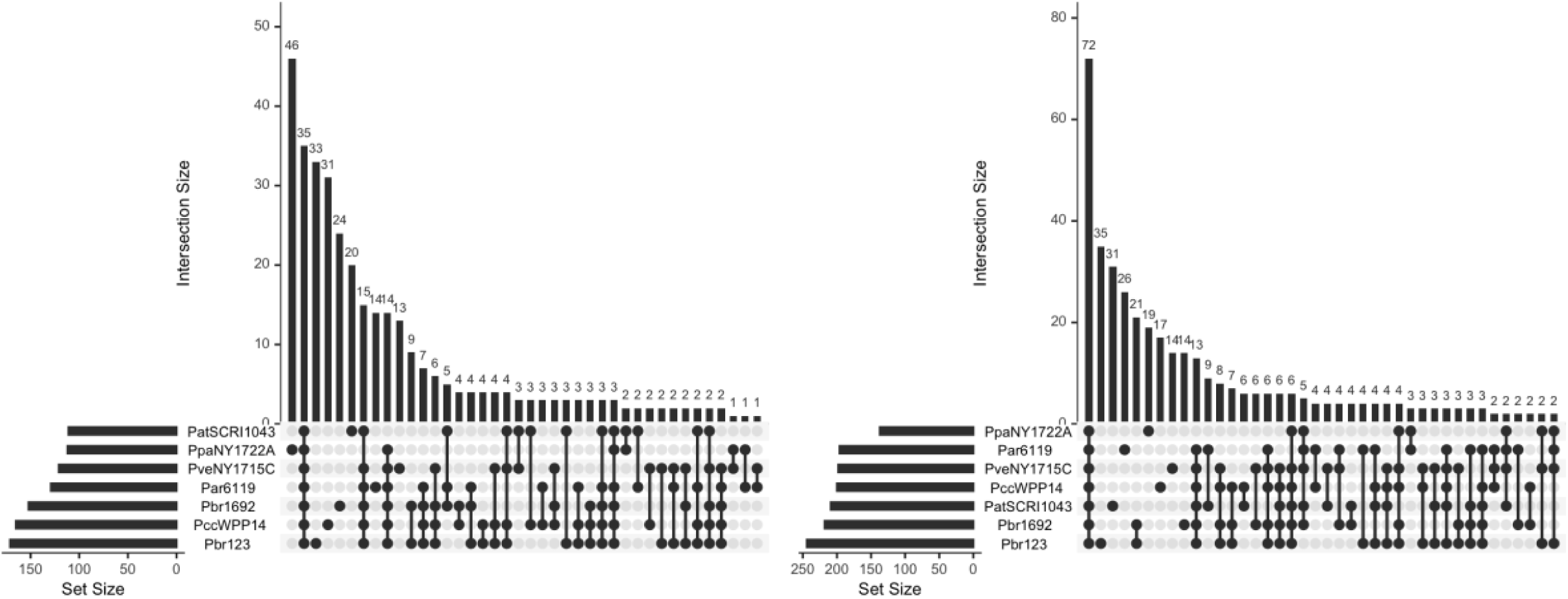
Shared orthogroups between strains, where the absolute(mean fitness) of the corresponding gene was >1 in M9 with glucose (left) or polygalacturonate (right) as a carbon source. UpSet plots were generated using “UpSetR” (Conway et al., 2017) in R.

### Several conserved Enterobacterial Common Antigen (ECA) genes were broadly important in minimal media and tubers

The outer membrane of strains in the order Enterobacterales, including *Pectobacterium*, contains a cell envelope-associated carbohydrate polymer known as the Enterobacterial Common Antigen (ECA) (Weckener et al., 2023). The highly-conserved biosynthetic gene cluster is made up of *wecA* (*rfe*), *wzzE*, *wecB* (*rffE*), *wecC* (*rffD*), *rffG*, *wecD* (*rffC*), *wecE* (*rffA*), *wzxE*, *wecF* (*rffT*), *wzyE*, and *wecG* (*rffM*); HER17_RS01145 through HER17_RS01195 in PccWPP14 (Hung et al., 2006; Li et al., 2018; Rai and Mitchell, 2020). Several genes in this cluster were important for growth in both media and plant conditions (Supplementary Figure 4). Specifically, *rffG* (HER17_RS01165) was essential in PccWPP14 and Par6119, had no data in Pbr123, Pbr1692, and PpaNY1722A due to a lack of insertions, and negative fitness values (<-2) across all conditions for PveNY1715C (Supplementary Figure 4). Interestingly, the fitness values for *rffG* (ECA_RS20810) in PatSCRI1043 appear to be condition-specific: for example, the mean fitness value in M9 with glucose = +1.8 compared to M9 with polygalacturonate = −4.1 (Supplementary Figure 4). Disruption phenotypes for *wzxE* and *wzyE* were also noteworthy. In Pbr123, *wzxE* (AC4LGO_01295) and *wzyE* (AC4LGO_01305) had strongly negative fitness values in most conditions tested. In PveNY1715C, *wzxE* (AC4LGM_01470) had negative fitness values in all conditions and no fitness data for *wzyE* (AC4LGM_01480). In PatSCRI1043, *wzxE* (ECA_RS20795) and *wzyE* (ECA_RS20785) had negative fitness (<-1) values in LB and tubers (Supplementary Figure 4).

### Core genes impacting fitness in potato tubers

Using the absolute(mean fitness) cutoff >1, there were 21 genes important for growth in potato tubers in all seven strains (Figure 5). Many of these metabolic genes, such as those involved in amino acid transport and metabolism, nucleotide transport and metabolism, and coenzyme transport and metabolism, were also broadly important for growth in minimal media, indicating a common conditional essentiality requirement for growth beyond optimal lab conditions. Due to the more variable *in vivo* experimental condition and generally weaker phenotypes observed, we also considered genes with absolute(mean fitness) <1 and >0.5. These cutoffs were applied to all genes in an orthogroup, and in many cases these genes had strong phenotypes in up to six strains tested, with a single strain weaker phenotype causing the orthogroup to not pass the more stringent filtering applied. Using this lower limit absolute(mean fitness) value added 15 additional genes to our list of orthogroups important in tubers. However, the other largest sets with up to 109 genes each highlight the strain-specific variability of this screen (Figure 5). Many genes important for growth in potato tubers were also important for growth in minimal media. Specifically, genes involved in nucleotide biosynthetic pathways (*carAB*, *purDH*, *pyrCD*) and the biosynthesis of various amino acids (*leuABCD*, *trpABCD*, *serA*) were conditionally essential in both minimal media and *in planta* conditions tested. The TonB system transport protein (HER17_RS10970) and TolC outer membrane channel (HER17_RS19825) were also important *in planta* and in most (though not all) *in vitro* conditions. This screen also identified the common catabolic genes specific to carbon sources available in tubers: *galK* (HER17_RS06390), *kdgK* (HER17_RS00410) and *kduD* (HER17_RS11410). These genes are conditionally essential for growth in polygalacturonate as a carbon source, with *galK* also being required for catabolism of galactose.

**Figure. 5.**
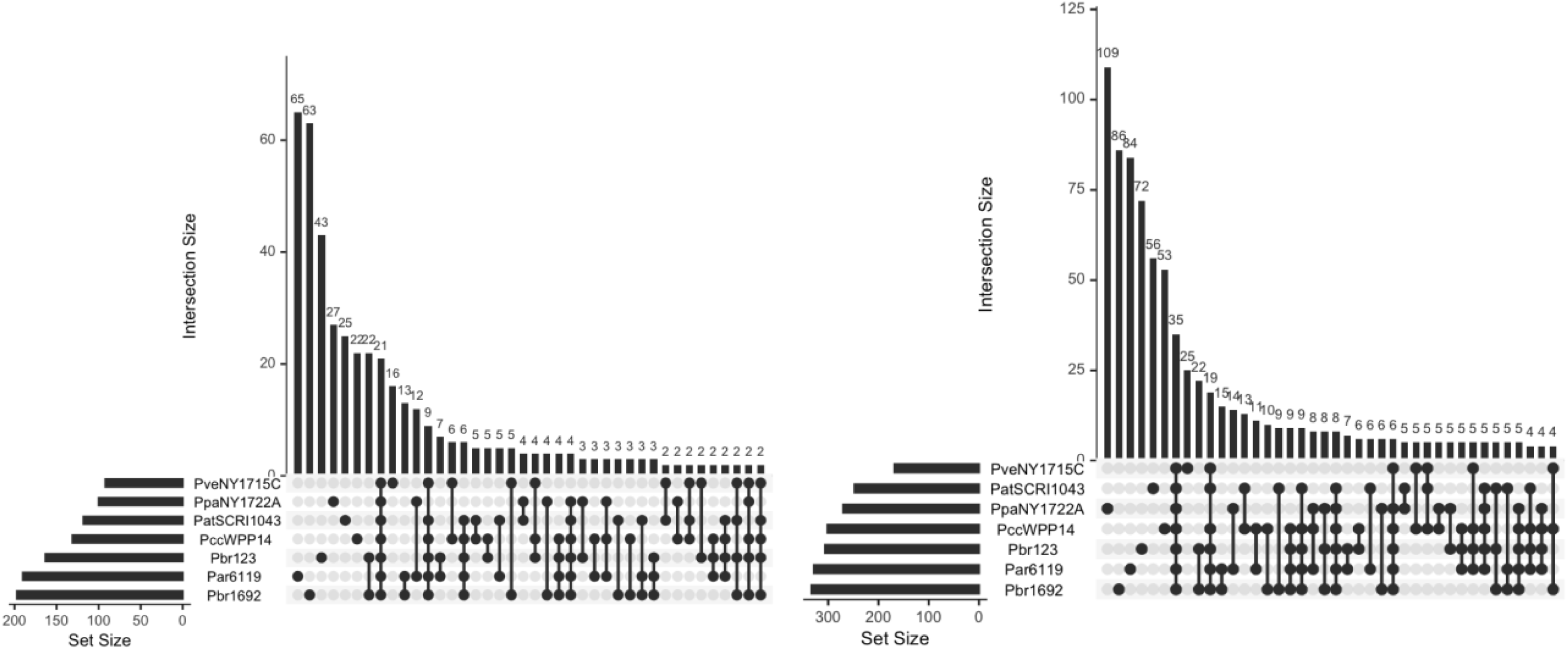
Shared orthogroups between strains, where the absolute(mean fitness) of the corresponding gene had fit >1 (left) and >0.5 (right) in potato tubers. UpSet plots were generated using “UpSetR” (Conway et al., 2017) in R.

### Putative T6SS toxin immunity proteins were conditionally essential in tubers

Interestingly, we identified an ankyrin repeat domain-containing protein encoding gene (HER17_RS04935) that was conditionally essential for growth of all strains in tubers (mean fit in tubers range −0.6 to −3.2), with no *in vitro* fitness phenotype. Since it is adjacent in all strains to a PAAR domain-containing protein and type VI secretion system (T6SS) tip protein VgrG, this ankyrin-domain containing protein is likely a type VI immunity protein, specifically important during growth in tubers. The orthologous gene in PatSCRI1043, ECA_RS16890 (old locus tag ECA3425), was previously noted to be present in a T6SS cluster (Arizala and Arif, 2019). Interestingly, while most strains tested here have a single copy of this gene at this location, PpaNY1722A has four adjacent copies and PatSCRI1043 has two adjacent gene copies. For all strains tested, only the first gene copy in each cluster has a fitness phenotype, despite the presence of paralogs in PpaNY1722A and PatSCRI1043 (Figure 6). The downstream co-localized DUF4123 domain-containing proteins in this cluster are reported molecular chaperones for the T6SS (Colautti et al., 2024).

**Figure 6:**
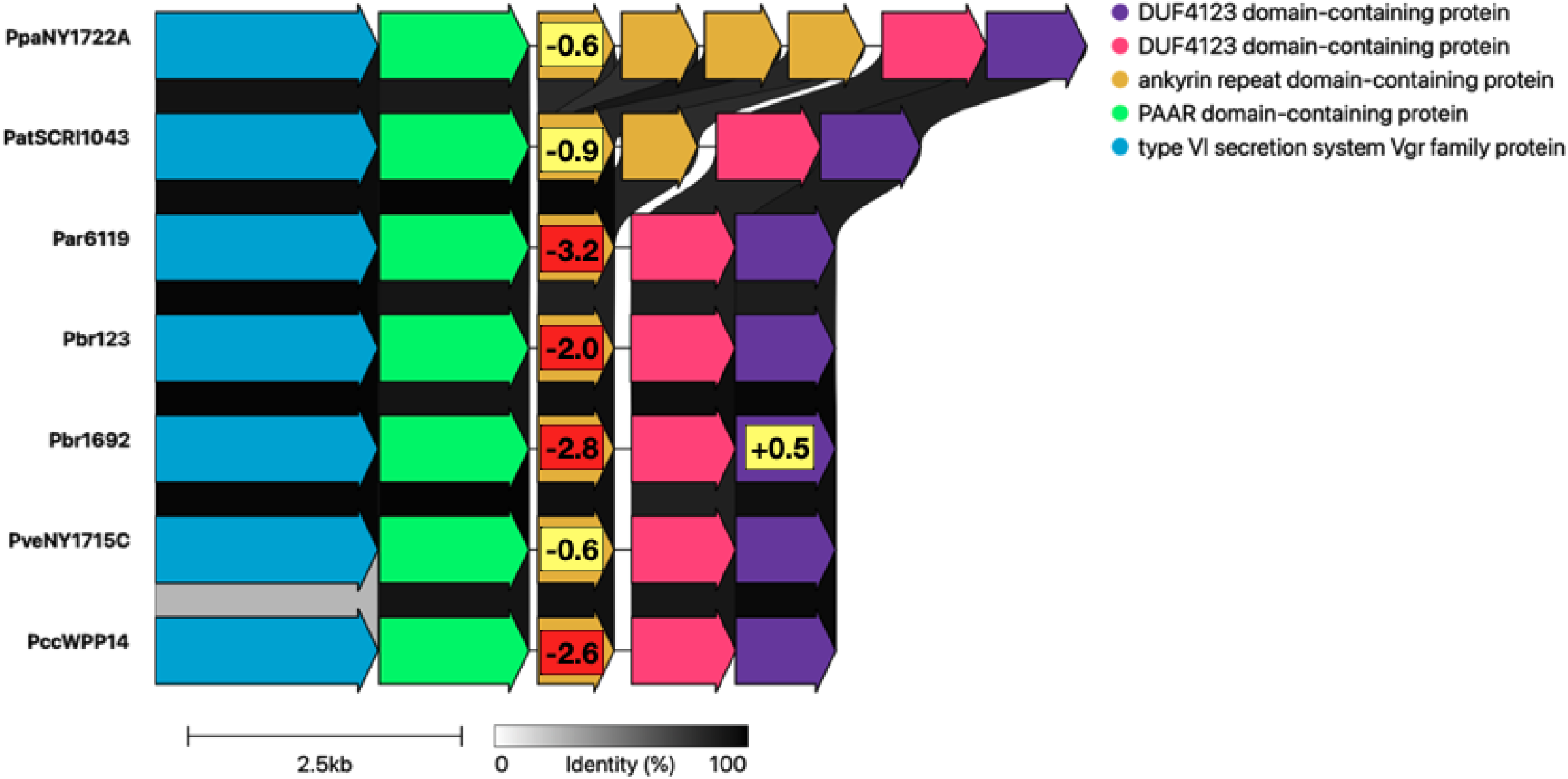
Ankyrin repeat domain-containing protein encoding genes commonly and specifically important in tubers, though phenotypes are limited to only the upstream copy when multiple copies exist. Gene synteny visualized using clinker (Gilchrist and Chooi, 2021) of the genomic regions immediately surrounding orthologous ankyrin repeat domain-contain protein encoding genes. Overlaid numbers are the mean fitness values for the tuber experimental conditions; where absolute(mean fitness) values <0.5 are not shown, values are highlighted in yellow (absolute(mean fitness) >0.5 and <2) or red (absolute(mean fitness) >2).

Similarly, we identified DUF1240 domain-containing protein encoding genes in similarly-organized genomic clusters, ranging from two to seven copies per cluster (Figure 7). Mean fitness values in tuber for the first gene in the cluster ranged from −0.7 to −3.0, with no data available (due to insufficient viable insertions and/or reads) for orthologous genes in Pbr123, PccWPP14, and PveNY1715C. Pbr123 was unique in that the second DUF1240 domain-containing protein encoding gene in the cluster (AC4LGO_03755) had a mean fitness in tubers of −0.7. DUF1240 has been previously proposed as an immunity protein family active against membrane disrupting type VI effectors (Fridman et al., 2020). Given the conserved and often large fitness defects of insertional mutants in the DUF1240 domain-containing proteins, we predict the upstream hypothetical proteins (HER17_RS03590) to be T6SS toxins.

**Figure 7:**
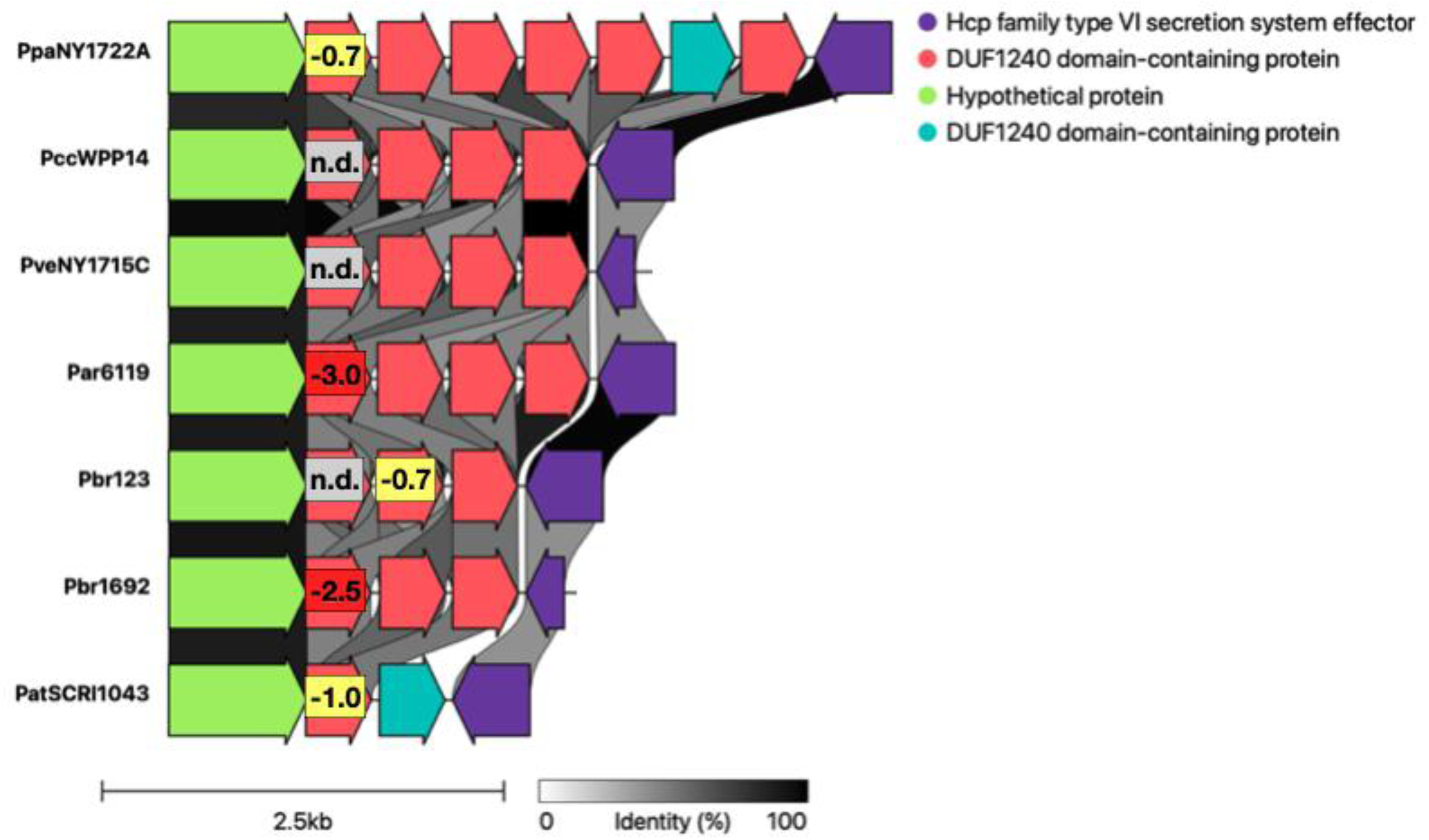
DUF1240 domain-containing protein encoding genes commonly and specifically important in tubers, though phenotypes are generally limited to only the upstream copy when multiple copies exist. Gene synteny visualized using clinker (Gilchrist and Chooi, 2021) of the genomic regions immediately surrounding orthologous DUF1240 domain-contain protein encoding genes. Overlaid numbers are the mean fitness values for the tuber experimental conditions; where absolute(mean fitness) values <0.5 are not shown, values are highlighted in yellow (absolute(mean fitness) >0.5 and <2) or red (absolute(mean fitness) >2). “N.d.” (grey) indicates “no data”, where insertions were present but in insufficient abundance to calculate fitness values for a given gene.

### Some putative toxin immunity proteins show strain-specific conditional essentiality

There were numerous genes that encode for putative hypothetical proteins that were specifically important in tubers and varied in either presence and/or disruption phenotype among the strains tested. Though the specific annotations varied (“hypothetical protein”, “membrane protein”, “DUF2645”, “DUF1240), genes encoding these proteins were often co-located with toxin-antitoxin genes or type VI secretion system effectors. For example, the protein of unknown function DUF2645 was present in only PccWPP14 (HER17_RS20130) and PveNY1715C (AC4LGM_21850) and gene disruptions had a large tuber-specific phenotype in these strains (mean fitness in tubers= −3.5 and −2.2). Both orthologs are located adjacent to Hcp family T6SS effectors (HER17_RS20135 and AC4LGM_21855), which were broadly dispensable when disrupted. Interestingly, the homologous DUF2645 family protein encoding gene in PccWPP14 (HER17_RS18025) did not have any phenotype when disrupted, suggesting that the adjacent Hcp family T6SS effector (HER17_RS18020) might not be activated under the conditions tested here. The DUF2645 encoding genes AC4LGN_02395 and AC4LGN_02400 in Par6119 also had a large contribution to competitive fitness in tubers and are downstream from the orthologous hypothetical and possibly toxin-encoding gene (mean fitness in tubers = −3.0 and −2.9) (Figure 8).

**Figure 8.**
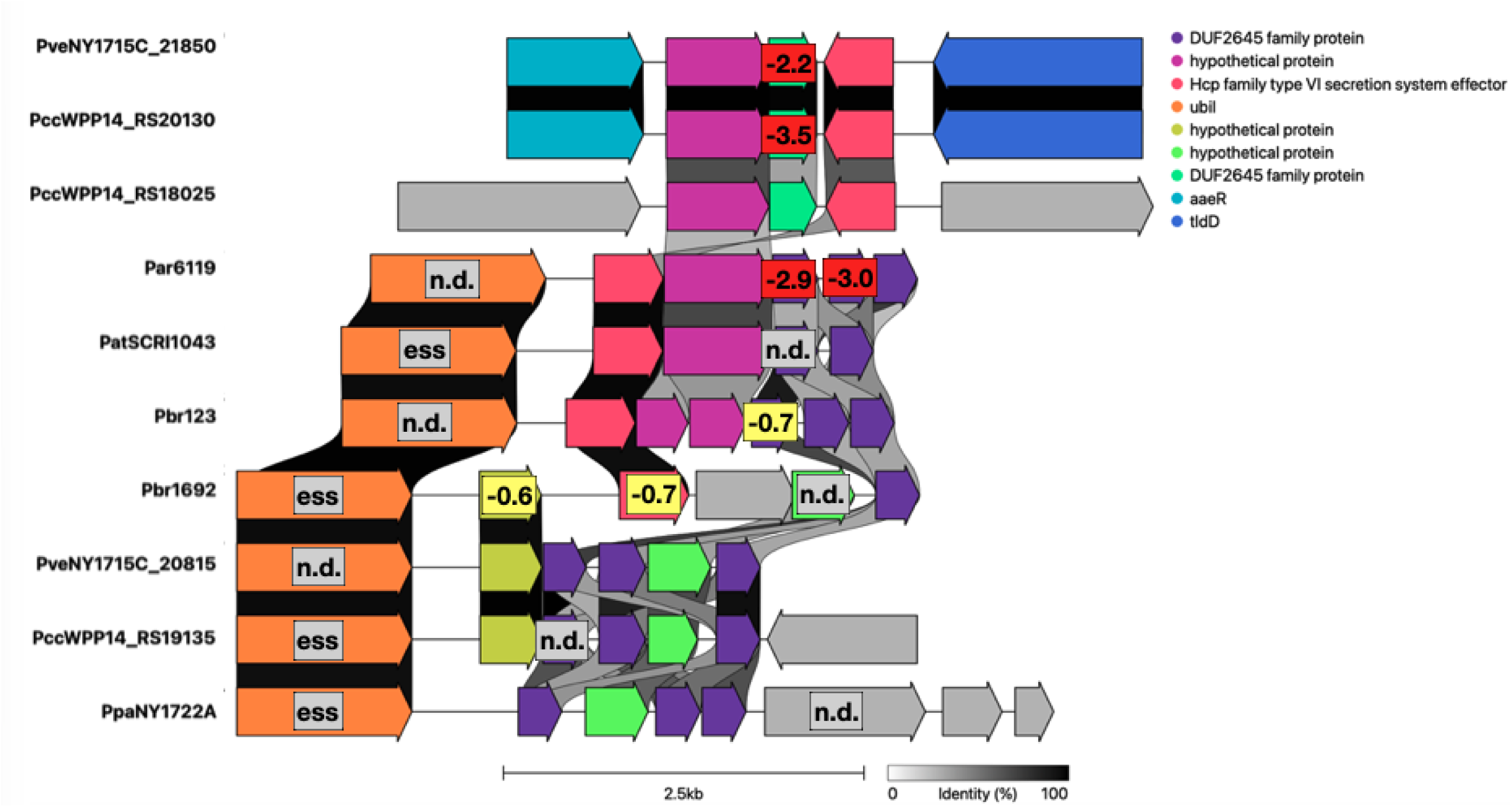
Genes encoding DUF2645 family proteins are likely type VI immunity genes. Gene synteny of the genomic regions immediately surrounding orthologous DUF2645 family protein encoding genes visualized using clinker (Gilchrist and Chooi, 2021). Overlaid numbers are the mean fitness values for the tuber experimental conditions, highlighted in yellow (mean fitness <-.5 and >-2) or red (mean fitness <-2). “N.d.” (grey) indicates “no data”, where insertions were present but in insufficient abundance to calculate fitness values for a given gene and “ess” (grey) indicates a likely essential gene based on the TnSeq data.

We also identified an Imm2 family immunity protein-coding gene specific to Par6119, *imm2* (AC4LGN_00350), with mean fitness in tubers = −2.1. This gene is downstream (on the negative strand) of AC4LGN_00355, encoding a DUF637 domain-containing protein with a predicted C-terminal toxin-deaminase domain. While Par6119 has a second adjacent copy of this Imm2 family immunity protein encoding gene (*imm2*, AC4LGN_00345), BarSeq phenotypes *in planta* were limited to AC4LGN_00350. The closest homologous genes on the Fitness Browser are in *Ralstonia solanacearum* (*R. solanacearum* UW163 UW163_RS10295 and *R. solanacearum* IBSBF1503 RALBFv3_RS03375) with 33% identity and no available insertion data (Georgoulis et al., 2021), implying they are likely essential in those strains.

### Strain-specific genes important for growth in tubers: carbapenem resistance

The antibiotic carbapenem is produced by several *Pectobacterium* and *Dickeya* species, with biosynthetic genes found in a single gene cluster also containing the genes required for carbapenem resistance (Shyntum et al., 2019). In this study, the biosynthetic cluster was present in both *P. brasiliense* strains tested, 1692 and 123-1. Interestingly, *P. parmentieri* have carbapenem resistance genes even though the biosynthesis genes are not present, noted previously in (Shyntum et al., 2019). This is apparent in the fitness results for the carbapenem immunity genes *carFG* (GT391_RS09030, GT391_RS09035) and an adjacent hypothetical protein encoding gene (GT391_RS09040), which are conditionally important in tubers for strains Pbr123 and Pbr1692, but not PpaNY1722 (Figure 9). Interestingly, mutants lacking conserved genes *mrcB* (penicillin-binding protein 1B, GT391_RS12785) and *lpoB* (penicillin-binding protein activator, GT391_RS19885) have comparable strong negative fitness phenotypes in tuber, specific to *P. brasiliense*.

**Figure 9:**
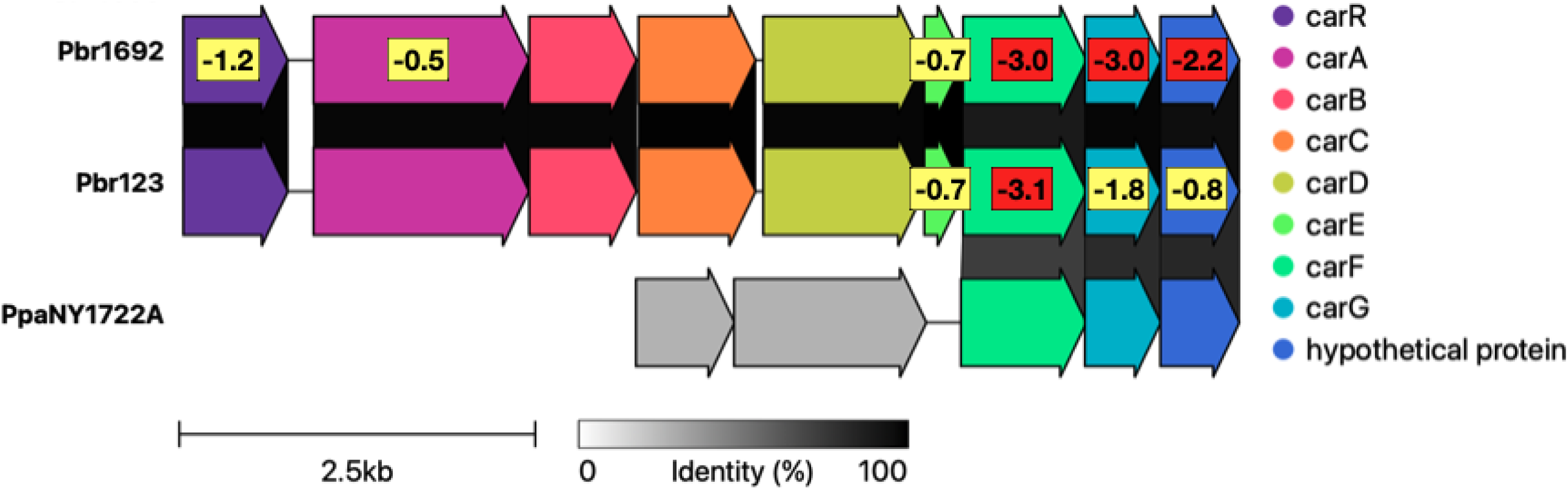
Carbapenem resistance genes *carFG* (GT391_RS09030, GT391_RS09035 in Pbr1692) and the adjacent hypothetical protein encoding gene (GT391_RS09040) present here only in Pbr123, Pbr1692, and PpaNY1722A, are important for full fitness only in carbapenem-producing *P. brasiliense* strains, and only in the tuber experiments. Gene synteny visualized using clinker (Gilchrist and Chooi, 2021). Overlaid numbers are the mean fitness values for the tuber experimental conditions, where absolute fitness values <0.5 are not shown, values are highlighted in yellow (absolute(mean fitness) >0.5 and <2) or red (absolute(mean fitness) >2).

### Strain-specific genes important for growth in tubers: LPS (*waa*) and group 4 capsule (G4C) O-antigen clusters

The *waa* cluster from *rfaD* (*waaD*) to *waaA* (in PccWPP14: HER17_RS20770 to HER17_RS20800), includes core lipopolysaccharide (LPS) biosynthesis genes (Ranjan et al., 2021). Enzymes encoded by this cluster are responsible for glycosidic linkages among the phosphorylated glycolipid (lipid A), core oligosaccharide, and surface-exposed glycan polymer (O-antigen) components of Pectobacterium LPS (Park et al., 2026). The beginning and end of this cluster are largely conserved, while the middle diverges among the seven strains analyzed here (Figure 10). The beginning of the cluster encompasses *rfaD* (HER17_RS20770), *rfaF* (HER17_RS20775), *rfaC* (HER17_RS20780), *rfaQ* (HER17_RS20785), and an O-antigen ligase family protein, *rfaL* (HER17_RS20790) (Park et al., 2026). Though *rfaD*, *rfaF*, *rfaC*, and *rfaQ* are conserved, fitness values vary. For example, fitness values for *rfaD* and *rfaF* ranged from nearly 0 (PveNY1715C) to strongly negative (Par6119) (Figure 10). In Par6119, disruption of *rfaD* (AC4LGN_21255), *rfaF* (AC4LGN_21260), and *rfaC* (AC4LGN_21265) had strong negative fitness effects specific to potato tubers (mean fitness in tubers = −2, −2.3, −2.1 respectively). In PccWPP14 and Pbr123, disruptions of these genes resulted in moderate phenotypes in tubers between −0.5 and −1.0 (Figure 10). There is high genomic diversity in the middle of the cluster (between *rfaC* and *waaA*). PccWPP14 and Pbr123 only have three genes (HER17_RS20785 through HER17_RS20795) while PpaNY1722A contains eight genes not found in the other six strains, of which five (AC4LGQ_22375 through AC4LGQ_22395) have tuber-specific negative fitness values ranging from −0.7 to −1.2 (Figure 10). Par6119 is also noteworthy in that the genes *waaG* (AC4LGN_21280), *waaO* (AC4LGN_21285), and AC4LGN_21305 encoding a glycosyltransferase family 2 protein have strong negative fitness values specific to tubers (mean fitness in tubers = −3.4, −3.1, and −3.5 respectively). In PveNY1715C, disruptions in a different glycosyltransferase family 2 protein (AC4LGM_22535) have a strong tuber-specific phenotype (mean tuber fitness = −2.7). The gene at the end of the cluster, *waaA* (HER17_RS20800) is universally either essential or lacking fitness data (Figure 10).

**Figure 10.**
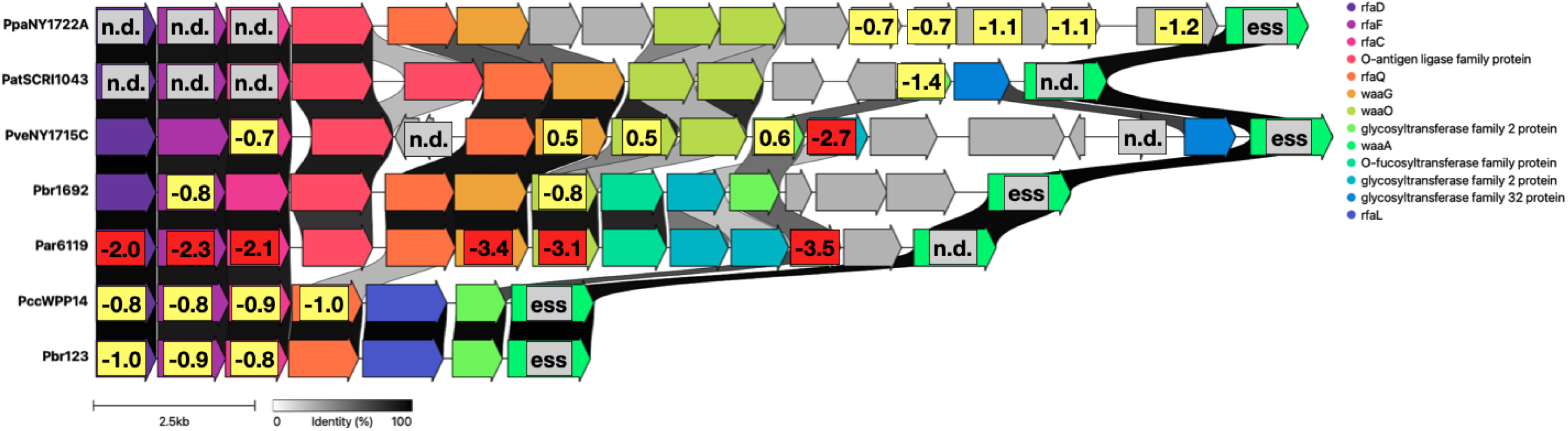
Gene synteny visualized using clinker (Gilchrist and Chooi, 2021) of the core LPS cluster, from *rfaD* to *waaA*, showing variation among these seven strains. While *waaA* is likely essential in all strains, there is a range in both cluster size and gene fitness values in tubers (for example, *waaG* and *waaO* in Par6119). Overlaid numbers are the mean fitness values for the tuber experimental conditions, highlighted in yellow (mean fitness <-.5 and >-2) or red (mean fitness <-2). “N.d.” (grey) indicates “no data”, where insertions were present but in insufficient abundance to calculate fitness values for a given gene and “ess” (grey) indicates a likely essential gene based on the TnSeq data.

The group 4 capsule (G4C) gene cluster encodes enzymes responsible for biosynthesis of surface protein-attached polysaccharides that are frequently identical to cognate LPS O-antigen. Therefore, the G4C has also been called the O-antigen capsule (Peleg et al., 2005; Ranjan et al., 2021). Park *et al*. identified enzymes within this cluster as responsible for O-antigen biosynthesis (Park et al., 2026). This gene cluster, defined as *wecA* (HER17_RS14615) to the YjbH domain-containing protein encoding gene (HER17_RS14485) (Ranjan et al., 2021; Park et al., 2026), varies in internal composition among the seven strains examined here (Supplementary Figure 5). Pbr123 is notable as the only strain with negative fitness values for almost all (24 of 27) genes in the cluster in most (but not all) M9 minimal media and tuber conditions (Figure 11). The genes *rbfA* (HER17_RS14595) and HER17_RS02220, a methionyl-tRNA formyltransferase, are essential in three and five of the seven strains, respectively. The gene *rfbA* is involved in biosynthesis of dTDP-rhamnose, which is a precursor for the biosynthesis of various cell wall polysaccharides, EPS, and LPS (Ranjan et al., 2021). Pbr1692 has two non-essential *rfbA* genes (GT391_RS00210 and GT391_RS00275). Notably, ECA_RS07135, encoding a glycosyltransferase family 2 protein, is essential in PatSCRI1043, where neither *rbfA* or the methionyl-tRNA formyltransferase are essential.

**Figure 11.**
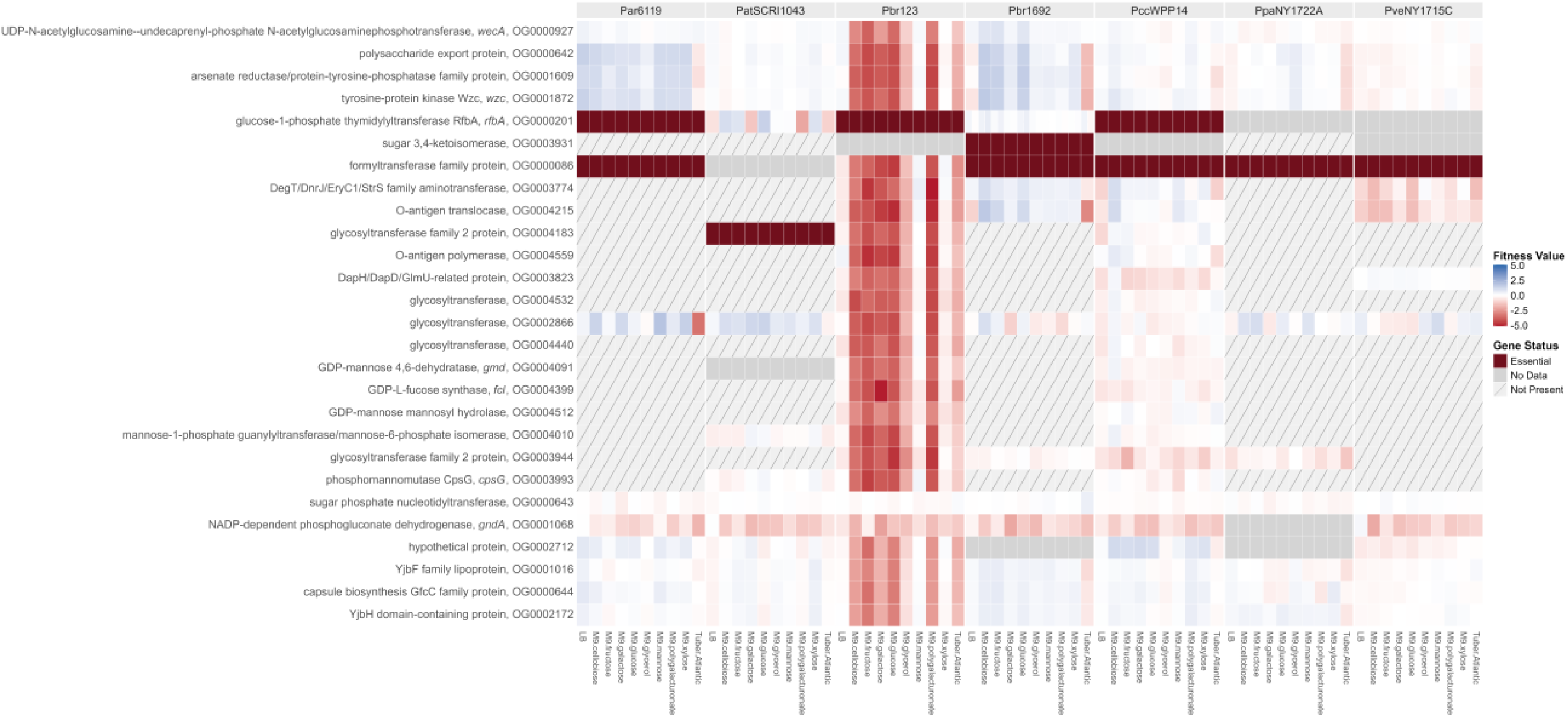
Mutations in the group 4 capsule (G4C) O-antigen cluster decrease fitness of Pbr123 across most M9 minimal media and tuber conditions. The G4C cluster is conserved between PccWPP14 and Pbr123 and this 27-gene cluster was selected to anchor the heatmap. Strain-specific genes not in the conserved PccWPP14 and Pbr123 cluster are not included. Synteny of the G4C cluster in all strains is shown in Supplementary Figure 5. Pbr1692 has two *rfbA* genes (GT391_RS00210 and GT391_RS00275), so the fitness tiles are divided. Heatmap was generated using “ggplot2” (Wickham, 2016) in R.

## Discussion

In this study, we aimed to leverage the natural genomic diversity of the *Pectobacterium* genus and the powerful high-throughput screening tool RB-TnSeq, to identify genes that are conditionally essential or important for bacterial growth (as competitive “fitness”) during soft rot infection of potato tubers. We purposefully included two strains, Par6119 and Pbr123, that were originally isolated from plant hosts other than potato yet still cause disease on potato tubers, to attempt to identify functional commonalities.

Furthermore, we tested these transposon libraries in several *in vitro* conditions including single sugars as carbon and energy sources that these pathogens would feasibly encounter in the potato tuber environment. A subset of these *Pectobacterium* species is capable of growth at elevated temperatures (i.e. >33°C); we generated heat shock fitness data for *P. brasiliense, P. carotovorum,* and *P. versatile.* These results were processed within this larger dataset, but a complete analysis will be presented in a separate work.

As part of the TnSeq mapping, it is possible to predict genes that are likely essential for growth in the rich medium LB, due to a lack of recovered insertion mutants. To the best of our knowledge, two groups have predicted *Pectobacterium* essential genes in rich medium using genome-wide metabolic modeling (Wang et al., 2015; Zoledowska et al., 2019). While we identified many of the same genes involved in core metabolism as being likely essential for growth in rich medium, these simulations significantly undercount total essential genes when considering non-metabolic genes. For example, our comprehensive mutagenesis and growth experiments identified essential non-metabolic genes such as those involved in ribosome formation, tRNA-ligases, DNA polymerase, and ATP synthases. Baltenneck et al. (Baltenneck et al., 2026) recently released a comparable dataset screening three *Pectobacterium* species by TnSeq to identify core essential genes, and their experimental results are more closely aligned with those described here.

In the context of the core genes of the *Pectobacterium* pangenome, we expected many commonalities between the species tested here, where important genes should be consistently identified in this screen across diverse genomic backgrounds. For example, the ability to produce PCWDEs and degrade pectin is a well-characterized virulence trait of the soft rot Pectobacteriaceae. Though the secreted pectate lyases were individually dispensable in this screen due to both redundancy and *in trans* complementation, genes involved in pectin and polygalacturonate catabolism (such as *kduD* and *kdgK*) were conditionally essential in the tuber experiments across all six species tested. Consistent with numerous previous TnSeq studies performed in the related plant pathogen *Dickeya* (Royet et al., 2019; Helmann et al., 2022; Robic et al., 2023), genes involved in the biosynthesis of amino acids and nucleotides were critical for bacterial growth both *in planta* as well as in minimal media conditions. To generate better context for the analysis of fitness factors for growth in potato tubers, we tested minimal media containing distinct carbon sources, some of which would be expected to be abundantly present in potato tubers (sucrose, glucose, fructose, galactose) and others that would be expected to be less abundant (cellobiose, mannose, xylose) (Knowles et al., 2009; Frost et al., 2016). The minimal medium condition with polygalacturonate as a carbon source was most similar to the tuber condition in terms of important metabolic genes. This suggests that while potato tubers contain a variety of available nutrients, the ability to break down and assimilate pectin is a very important trait for sustained bacterial population growth and generation of visible disease symptoms. Broadly speaking, genes in the COG categories involved in the biosynthesis of amino acids, nucleotides, and cofactors were most commonly important during growth in both minimal media and potato tubers.

There were many genes known to be important for tuber colonization that were not identified in this screen as a result of our methods and experimental design. It is important to note that well-characterized and critically-important secreted virulence factors such as plant cell wall degrading enzymes (PCWDEs) are not well suited to be identified in a mixed-population screen such as this. We also did not see any phenotypes for genes involved any secretion systems (type I, II, III, VI). This is likely due to the complementation that occurs when individual deficient mutants are co-localized in a larger population with wild type (at that locus) cells.

In some cases we observed differences in the impact of the disruption of a gene in whether disruption resulted in increased or decreased fitness. For example, it was interesting that the phenotypic direction for the *slyA* mutants differed between negative in Pbr1692 and positive for Pbr123 (and others). The SlyA regulon is a complex network and linked with PCWDEs, carbapenem biosynthetic genes, and PhoP (Bellieny-Rabelo et al., 2020). While we did not test virulence directly, our results showing decreased competitive growth for the Pbr1692 *slyA* mutant in tubers are consistent with this previous result. However, the observation that in other strains, such as Pbr123, the loss of *slyA* appears to increase growth should be further explored. It is possible that the SlyA regulon differs between the strains. It is also possible that in Pbr123 and other strains, *slyA* is functionally complemented *in trans* by wild-type levels of PCWDEs produced by neighboring cells and/or cell wall degradation products during the infection process. According to a recent analysis by Jonkheer et al., *P. brasiliense* has the highest genomic diversity of all *Pectobacterium* species (Jonkheer et al., 2021). Therefore, it is possible that broader genomic or regulatory differences between Pbr1692 and Pbr123 result in the divergence of *slyA* phenotypes seen here.

We examined the contributions to competitive fitness of three gene clusters contributing to the structure of the outer cell membrane: the Enterobacterial Common Antigen (ECA) cluster, the *waa* (LPS) cluster, and the group 4 capsule (G4C) or O-antigen capsule cluster. The ECA is a highly-conserved surface antigen present in nearly all members of Enterobacterales. Its invariability is noteworthy considering the order contains both plant and animal pathogens. In the past, study of ECA biosynthesis has been complicated, in part, due to the extensive interactions with O-antigen and peptidoglycan biosynthesis pathways (Rai and Mitchell, 2020). Recent work in models such as *Escherichia* and *Salmonella* has clarified protein function to some extent (Bennett and Mitchell, 2025), yet the biological significance of the ECA in each genus, remains largely unknown (Rai and Mitchell, 2020). In *Pectobacterium* spp., the ECA has been described as a pathogenicity determinant (Li et al., 2018). In our study, the ECA appears more broadly important for growth, with *rffG* predicted to be essential in Par6119 and PccWPP14 and has no fitness data due to a lack of insertions in Pbr123, Pbr1692, and PpaNY1722A (Supplementary Figure 4). A screen by Toth et al. isolated a bacteriophage-resistant *rffG* mutant in *P. atrosepticum* SCRI1043 with cell surface defects and reduced synthesis of the pectate lyase and cellulase virulence factors (Toth et al., 1999). The genes *wzxE* and *wzyE* were conditionally-essential or lacking data in a strain-specific manner, suggesting potential variability in biological significance (Supplementary Figure 4). Collectively, our results indicate that ECA biosynthesis can be broadly important for bacterial growth and competitive fitness, although the contribution of individual pathway components varies among strains.

The *waa* cluster is responsible for core LPS biosynthesis (Ranjan et al., 2021). Genes at the beginning of the cluster (*rfaD*, *rfaF*, *rfaC*, *rfaQ* and *rfaL*) were recently characterized in *P. carotovorum* PCC27 as contributing to virulence on kimchi cabbage (Park et al., 2026). Additionally, mutants in *rfaD*, *rfaF* and *rfaC* exhibited reduced proteolytic activity and increased membrane permeability, and motility was also reduced in *rfaD* and *rfaC* mutants (Park et al., 2026). Here, we were unable to generate fitness data for *rfaD*, *rfaF*, or *rfaC* in PatSCRI1043 or PpaNY1722A. We observed tuber-specific strong (<-2) fitness values in Par6119 and moderate (−0.7 to −1.0) in one to three of these genes in all other strains. It should be noted that Park et al. also attempted to generate a *kdtA* (*waaA*) mutant, but were unsuccessful and reported this may suggest the gene is essential (Park et al., 2026). Our analysis classified *waaA* as essential (or lacking data) in all tested strains (Figure 10). The results of Park et al. suggest that mutants may have additional phenotypes (unrelated to growth), such as decreased virulence and motility (Park et al., 2026). These findings highlight the importance of core LPS biosynthesis for fitness *in planta*, and suggest that the effects of *waa* mutations on competitive fitness may reflect alterations in multiple physiological processes, rather than growth defects alone.

The G4C or O-antigen cluster was highly variable between these seven strains, both in cluster organization and gene contributions to competitive fitness. Conservation of 27 genes in this cluster between PccWPP14 and Pbr123 (Supplementary Figure 5) allowed us to anchor our analysis, focusing on those genes that are present in at least those two strains. Interestingly, the loss of nearly every gene in Pbr123 has a strong negative fitness effect, particularly in minimal media conditions (Figure 11). These results were not observed with disruptions to orthologous genes in PccWPP14, suggesting strain-specific differences in the relative contribution to growth of this conserved structural component.

In the experimental conditions tested here, the induction of intercellular warfare strategies like the production of T6SS effectors and the antibiotic carbapenem were specifically observed in the tubers. Induction signals for the T6SS and carbapenem production are complex, and include quorum sensing and other environmental signals (McGowan et al., 1995; Liu et al., 2008). Experiments performed here highlighted the important systems specifically in potato tubers during soft rot infection of type VI immunity proteins, carbapenem resistance genes in *P. brasiliense*, and hypothetical proteins that are likely antitoxin components of toxin-antitoxin systems. Mutagenesis screens are a highly-effective method to identify essential or conditionally essential loss-of-function mutants; with examples seen frequently here in putative antitoxin and type VI immunity genes. Baltenneck et al. also highlighted strain-specific toxin-antitoxin and contact-dependent growth inhibition (CDI) systems as having diverse essentiality profiles (Baltenneck et al., 2026). Some antitoxin genes in PatSCRI1043 were identified as essential in our study as well as previously, such as the helix-turn-helix domain-containing proteins ECA_RS03355 (ECA0673) and ECA_RS14305 (ECA2911) (Baltenneck et al., 2026).

By adding the functional genomic screening data across these six *Pectobacterium* species to the two *Dickeya* species (Helmann et al., 2022; Gonzalez-Tobon et al., 2024) on the interactive Fitness Browser (https://fit.genomics.lbl.gov/) and its broader data ecosystem (Price and Arkin, 2024), we have continued to build a foundational dataset for the soft rot Pectobacteriaceae. We also specifically generated transposon mutant libraries in two *P. brasiliense* strains for comparative analysis between these strains with different hosts of isolation. This resource will be valuable for those undertaking analysis of additional functional data for these and related strains within the broader genomic context of these agriculturally-important soft rot pathogens.

## Materials and Methods

### Bacterial strains and growth conditions

Strains used in this study are described in Supplementary Table 6. All bacterial strains were routinely cultured in LB medium (per L: 10 g tryptone, 5 g yeast extract, and 10 g NaCl) with *Pectobacterium* spp. at 28°C, and *E. coli* at 37°C. When noted, kanamycin was used at a final concentration of 50 μg/ml. For minimal medium fitness assays, strains were grown in M9 minimal medium (M9) as described in [https://cshprotocols.cshlp.org/content/2006/1/pdb.rec614] with 0.4% (w/v) carbon source (one of): cellobiose, fructose, galactose, glucose, glycerol, mannose, myo-inositol, polygalacturonate, starch, sucrose, or xylose.

### Whole-genome sequencing and genome annotation

High-quality genomes were developed for four of the strains used in this study: *P. aroidearum* 61-19 (Par6119), *P. brasiliense* 123-1 (Pbr123), *P. parmentieri* NY1722A (PpaNY1722A), and *P. versatile* NY1715C (PveNY1715C). For Par6119 and Pbr123, high-molecular-weight genomic DNA was extracted with the Monarch HMW DNA Extraction Kit (New England Biolabs, United States) following the protocol for bacteria. DNA quantity was assessed using a Qubit 3.0 fluorometer (Thermo Fisher Scientific, United States), and DNA integrity was assessed by agarose gel electrophoresis. The isolated DNA was used to prepare a Nanopore sequencing library using the ONT Rapid Barcoding Sequencing kit (SQK-RBK004) (Oxford Nanopore Technologies, U.K.). The sequencing reaction was run in a MinION flow cell. Base calling was performed using Guppy (Oxford Nanopore Technologies, U.K.) and raw reads were de-multiplexed with QCAT (https://github.com/nanoporetech/qcat). Supplementary Table 7 details Nanopore coverage results. The isolated DNA was also used to prepare an Illumina sequencing library using the Nextera DNA Flex Library Prep Kit (Illumina Inc., United States). The DNA library was sequenced on a NextSeq 500 (Illumina Inc., United States) by the Cornell Genomics Facility.

Bacterial genome sequencing for *P. parmentieri* NY1722A and *P. versatile* NY1715C was performed by Plasmidsaurus using Oxford Nanopore and Illumina Sequencing Technology (“Big Hybrid”) with custom analysis and annotation. The exact method used to assemble the Nanopore and Illumina reads into complete genomes varied from strain to strain. In short, consensus assemblies were constructed using either Trycycler (Wick et al., 2021) or Autocycler (Wick et al., 2025). The strain-specific details, including software versions, can be found in the metadata for the corresponding NCBI GenBank records. Generally, the following method was used to assemble the genomes: the raw Nanopore reads were filtered and trimmed using Filtlong (https://github.com/rrwick/Filtlong) using parameters “--min_length 1000 --keep_percent 95”. The raw Illumina reads were filtered and trimmed using Fastp (Chen et al., 2018). Subsets of the filtered Nanopore reads were constructed using the “trycycler subsample” or the “autocycler subsample” commands. Rough assemblies were constructed from each of these subsets using Flye (Kolmogorov et al., 2019), Minism (Li, 2016), and Raven (Vaser and Sikic, 2021). Consensus assemblies were constructed from the rough assemblies using the standard sequence of commands (either https://github.com/rrwick/Trycycler/wiki/How-to-run-Trycycler or https://github.com/rrwick/Autocycler/wiki/Fully-automated-assembly). The generated consensus assembly was polished with the filtered Nanopore reads using Medaka (https://github.com/nanoporetech/medaka) and with the filtered Illumina reads using Nextpolish (Hu et al., 2020) or Polypolish (Wick and Holt, 2022). The correctness of the assembly was validated by constructing a second, independent assembly with Unicycler (Wick et al., 2017). The taxon of the final assembly was verified using NCBI’s prokaryotic genome annotation pipeline (PGAP) (Tatusova et al., 2016) or ReferenceSeeker (Schwengers et al., 2020). The completeness of the assembly was checked using BUSCO (Seppey et al., 2019) using the “pectobacterium” lineage database. All genomes were uploaded to NCBI and annotated using NCBI’s PGAP.

### Identification of orthogroups

All “complete”- and “chromosome”-level *Pectobacterium* genome assemblies (N=152) were downloaded from NCBI using NCBI Datasets command line tools (O’Leary et al., 2024) with parameters “taxon Pectobacterium --assembly-level complete,chromosome --include protein” on 7/7/2026. These assemblies were combined with the closed assemblies for the strains *P. aroidearum* 61-19, *P. brasiliense* 123-1, *P. parmentieri* NY1722A, and *P. versatile* NY1715C. We used the CheckM v1.2.4 (Parks et al., 2015) lineage workflow to calculate genome completeness and contamination, with “Enterobacteriaceae” marker genes. Based on these results, we removed the outlier genome *Pectobacterium* sp A5354 (completeness = 90%) from analysis. All remaining genomes (N=151 NCBI + 4 additional) had completeness >94% and contamination ≤1%. Orthologous genes were identified from the complete set of amino acid FASTA files and assigned to Orthogroups (OGs) using OrthoFinder v3.1.3 (Emms and Kelly, 2019). We combined the OrthoFinder output files “Orthogroups.tsv” and “Orthogroups_UnassignedGenes.tsv” to ensure all genes, including strain-specific genes, were associated with OG numbers for downstream analysis. For each strain, all remaining genomic features present in the TnSeq or BarSeq data that were not included as OrthoFinder inputs due to lacking protein sequence (ie. tRNAs, pseudogenes) were assigned to a singleton orthogroup with their locusId as its identifier. All gene synteny visualizations were generated using clinker (Gilchrist and Chooi, 2021).

### Construction of barcoded transposon libraries

Barcoded mutant libraries were constructed by conjugating barcoded transposon plasmids from the *E. coli* WM3064 donor libraries: *mariner* transposon plasmid pKMW3 (APA752) or Tn5 transposon plasmid pKMW7 (APA766) (Wetmore et al., 2015) into the wild-type strains, as in (Helmann et al., 2022). Briefly, donor and recipient strain mixtures were incubated for one day on LB supplemented with diaminopimelic acid (DAP) at 28°C. Exconjugants were selected using LB kanamycin at incubated at 28°C for 1 day. All colonies were resuspended in LB with kanamycin, diluted to a starting OD600 of 0.2 in 200 ml LB kanamycin and grown at 28°C with shaking for 4 hours to a OD600 ∼ 1.5. Glycerol was added to the library to a final concentration of 15% and 1ml aliquots were frozen at −80°C.

### RB-TnSeq mapping and identification of likely essential genes

For transposon insertion-site sequencing (TnSeq), multiple 1ml aliquots of each library were pelleted, and genomic DNA was purified using the Monarch Genomic DNA Purification Kit (New England Biolabs, United States). Samples were eluted in 100µl elution buffer and quantified using a NanoDrop One (Thermo Fischer Scientific, United States). For each library, we sequenced 3-6 independent freezer aliquots, starting with 500ng genomic DNA as input for library preparation using the NEBnext Ultra II FS DNA Library Prep kit (New England Biolabs, United States), following manufacturer’s instructions with some modifications detailed previously (Helmann et al., 2022). We used a modified version of the protocol initially detailed in (Wetmore et al., 2015), with a modified two-step PCR based on (Rubin et al., 2022). DNA libraries were submitted for 1×150bp sequencing (Mid-Output Kit) at the Biotechnology Resource Center (BRC) Genomics Facility at the Cornell Institute of Biotechnology sequencing on a NextSeq 500/550 (Illumina, Inc., United States).

Sequence data were analyzed using the FEBA scripts (Wetmore et al., 2015). For each strain, SetupOrg.pl was used to prepare the input genomic files starting from the GenBank (.gbk) DNA sequence and annotation file. We used MapTnSeq.pl to identify the barcode sequence and genomic location of each read [with identifiable transposon sequence], based on the “model_pKMW3.2” (*mariner* libraries) or “model_pKMW7.2” (Tn5 libraries) sequence. Finally, DesignRandomPool.pl was used to assemble the mutant pool using barcodes seen in a single location 10 or more times. Based on the TnSeq mapping data and insertion frequency per gene, we calculated likely essential genes using Essentiality.pl and the “Essentials” function from lib/comb.R (Wetmore et al., 2015). Protein-coding genes are considered to be essential or nearly essential if the normalized central insertion density score and normalized read density are <0.2 (Price et al., 2018). For these libraries, genes shorter than 100-225bp were excluded from analysis.

### Fitness experiments

For each genome-wide gene fitness experiment, an entire library freezer aliquot was thawed and recovered in 25 ml LB kanamycin at 28°C until OD600 1.0-1.5; approximate 6 hours. At this point, two 1 ml cell pellets were collected and frozen at −20°C as time0 measurements, and the remaining culture was washed in 10 mM KPO_4_ and used to inoculate experimental samples. All *in vitro* experiments were performed in 24-well cell culture plates, with 50µl standardized culture inoculated into 950µl media, with shaking overnight at 28°C. Experiments to test heat stress were performed in LB with shaking overnight at 36°C. After culture growth, the entire 1ml for each replicate was collected as a cell pellet, supernatant removed, and frozen at −20°C before gDNA purification.

For *in planta* experiments, *Solanum tuberosum* cv. Atlantic tubers were rinsed and surface sterilized using 70% ethanol. Inoculum was standardized to 10^9^ CFU/ml and 10µl was infiltrated into two replicate stab wounds per tuber created by pushing a 200µl pipette tip ∼3mm into the tuber. Six replicate tubers were inoculated for each transposon library. Tubers were sealed in clear plastic bags and stored in the dark at 28°C for 2 days. Bacterial cells were recovered from the sites of infection using a 1cm diameter cork borer to core the tubers. Duplicate samples from each tuber were combined in 8ml 10mM KPO_4_ and shaken at 200 rpm 28°C for 10 minutes. For each sample, 2ml of bacterial suspension was pelleted and frozen at −20°C before gDNA purification.

### BarSeq PCR and sequencing

Genomic DNA was purified from cell pellets using the Monarch Genomic DNA Purification Kit (New England Biolabs, United States). Purified DNA samples were eluted in 100µl elution buffer and quantified using a NanoDrop One. After purification, the 98°C BarSeq PCR as described in (Wetmore et al., 2015) was used to amplify the barcode region of each sample; in 50 μl total volume: 0.5 μl Q5 High-Fidelity DNA polymerase (New England Biolabs, United States), 10 μl 5X Q5 buffer, 10 μl 5X GC enhancer, 1 μl 10 mM dNTPs, 200-300 ng template gDNA, 2.5 μl common reverse primer (BarSeq_P1), and 2.5 μl of forward primer from one of the 96 indexed forward primers (BarSeq_P2_ITXXX), both at 10 μM (Wetmore et al., 2015). Following the BarSeq PCR, 10 μl of each reaction was pooled (up to 96 samples per pool), and 200 μl of this DNA pool was subsampled and purified using the DNA Clean and Concentrator Kit (Zymo Research, United States). The final DNA sequencing library was eluted in 30 μl nuclease-free water, quantified on a Nanodrop One, and submitted for sequencing at the BRC Genomics Facility at the Cornell Institute of Biotechnology. Prior to sequencing, the quality of each amplicon pool was assessed using a Bioanalyzer, and Pippin DNA size selection was used when needed to remove primers and over-amplification products. Each sequencing pool was submitted for a single 1×75bp sequencing run (High-Output Kit) on the NextSeq 500/550 (Illumina, Inc., United States).

### Calculation of fitness values

Sequencing data were used to calculate genome-wide gene fitness using the FEBA scripts originally published in (Wetmore et al., 2015). Scripts to calculate gene fitness values were run on a Cornell University BioHPC Cloud 60-core Linux (CentOS 7.6) server with 256 GB RAM. From the FASTQ reads for each sample, individual barcode sequences were identified and counted using MultiCodes.pl, and these counts were combined across samples for a given transposon library using combineBarSeq.pl. Using BarSeqR.pl, fitness values for each insertion strain were calculated as the log2 ratio of barcode abundance following library growth under a given experimental condition divided by the abundance in the time0 sample. The fitness of each gene is the weighted average of strain fitness values based on the “central” transposon insertions only, i.e., those within the central 10–90% of a gene. Barcode counts were summed between replicate time0 samples. For analysis, genes were required to have at least 30 reads per gene in the time0 sample, and 3 reads per individual strain (Wetmore et al., 2015). Gene fitness values were normalized across the chromosome so that the median gene fitness value was 0. All experiments described here passed previously described quality control metrics (Wetmore et al., 2015; Price et al., 2018). Fitness scores were plotted using “ggplot2” (Wickham, 2016) in R.

## Acknowledgments

We thank Bryan Swingle and Xing Ma for providing the *P. parmentieri* NY1722A and *P. versatile* NY1715C strains. We also thank Morgan Price for assistance with the FEBA analysis pipeline and data upload to the Fitness Browser. Sequencing was performed by the Biotechnology Resource Center (BRC) Genomics Facility (RRID:SCR_021727) at the Cornell Institute of Biotechnology. Mention of trade names or commercial products in this publication is solely for the purpose of providing specific information and does not imply recommendation or endorsement by the United States Department of Agriculture. USDA is an equal opportunity provider and employer.

## Data Availability

Genome assemblies for the newly sequenced strains and raw reads for whole-genome sequencing were submitted to NCBI under the BioProject PRJNA1471679, with sample metadata in Supplementary Table 8. Assembled genomes are as follows: *P. aroidearum* 61-19 (SAMN60462425), *P. brasiliense* 123-1 (SAMN60462424), *P. parmentieri* NY1722A (SAMN60462427), and *P. versatile* NY1715C (SAMN60462426). RB-TnSeq mapping reads (SUB16442135; Supplementary Table 9) and BarSeq reads (SUB16444671; Supplementary Table 10) were also submitted to NCBI under the BioProject PRJNA1471679. The analysis code for barcode identification, insertion-site mapping, essentiality prediction, and per-gene fitness analysis was previously published in (Wetmore et al., 2015) and is available at https://bitbucket.org/berkeleylab/feba/. Experimental metadata, genomic information and fitness values from BarSeq experiments that passed quality control metrics are available for all strains on the interactive Fitness Browser (https://fit.genomics.lbl.gov). Scripts used here for sequence data processing and analysis are available at https://github.com/tylerhelmann/2026-pectobacterium-RBTnSeq.

## Footnotes

1 As orthogroup identifiers are specific to this dataset, we have elected to use locus tags from PccWPP14 to identify genes within an orthogroup. Supplementary Table 1 can be used to determine corresponding locus tags in other strains.

