## Supplementary Figure for "Comparative functional genomics of six *Pectobacterium* species reveals core and unique fitness determinants in potato tubers"

### Supplemental Information

Supplementary Table 1. Full list of all orthogroups calculated using OrthoFinder v3. Separate tabs are included for OrthoFinder results files Orthogroups.tsv and Orthogroups_UnassignedGenes.tsv.

Supplementary Table 2. Essential gene predictions for the seven transposon libraries generated in this study. Data for each strain are included as separate tabs within this Excel document. Column "ess" indicates if a gene is a likely to be essential based on a lack of central transposon insertions. A note on the Pbr1692 gene list: the genome annotation (GCF_009873295.1) starts (i.e. coordinate position 1) in the middle of a gene (GT391_RS00005, encoding a non-ribosomal peptide synthetase). Because the gene "beginning" position (4,844,040) > "end" position (13,195), the FEBA pipeline drops this gene from analysis.

Supplementary Table 3. Number of replicate experiments for each strain and condition that passed previously-established QC metrics (Wetmore et al., 2015) and were uploaded to the Fitness Browser (https://fit.genomics.lbl.gov/).

| | Par6119 | PatSCRI1043 | Pbr123 | Pbr1692 | PccWPP14 | PpaNY1722A | PveNY1715C |
|---|---|---|---|---|---|---|---|
| LB | 3 | 3 | 5 | 5 | 11 | 4 | 5 |
| LB at 36˚C | 0 | 0 | 3 | 2 | 3 | 0 | 3 |
| M9+ cellobiose | 3 | 3 | 3 | 3 | 3 | 3 | 3 |
| M9+ fructose | 3 | 3 | 3 | 3 | 3 | 3 | 3 |
| M9+ galactose | 3 | 3 | 3 | 3 | 3 | 3 | 3 |
| M9+ glucose | 3 | 3 | 3 | 3 | 3 | 2 | 3 |
| M9+ glycerol | 3 | 3 | 3 | 3 | 8 | 3 | 3 |
| M9+ mannose | 3 | 3 | 3 | 3 | 3 | 3 | 3 |
| M9+ myo-inositol | 0 | 0 | 3 | 3 | 0 | 0 | 0 |
| M9+ polygalacturonate | 3 | 3 | 3 | 0 | 6 | 3 | 3 |
| M9+ starch | 0 | 0 | 0 | 0 | 3 | 0 | 0 |
| M9+ sucrose | 0 | 0 | 0 | 0 | 3 | 0 | 0 |

| | Par6119 | PatSCRI1043 | Pbr123 | Pbr1692 | PccWPP14 | PpaNY1722A | PveNY1715C |
|---|---|---|---|---|---|---|---|
| M9+ xylose | 3 | 3 | 3 | 3 | 3 | 3 | 3 |
| Tuber-Atlantic | 6 | 6 | 5 | 4 | 9 | 4 | 6 |

Supplementary Table 4. Lists of genes with strong phenotypes for each experiment. A strong phenotype is defined as |fit| > 2 and |t| > 5 (Wetmore et al., 2015). Data for each strain are included as separate tabs within this Excel document. Column “lrn” is “normalized gene fitness”.

Supplementary Table 5. Mean fitness for each gene under each condition tested. Data for each strain are included as separate tabs within this Excel document.

Supplementary Table 6. Bacterial strains used in this study.

| Strain | Description | Source |
|---|---|---|
| *Pectobacterium aroidearum* 61-19 | Wild-type strain, isolated from calla lily in NY, 2019 | This work |
| *Pectobacterium atrosepticum* SCRI1043 | Wild-type strain, isolated from potato in Scotland, 1985 | (Hinton et al., 1989) |
| *Pectobacterium brasiliense* 123-1 | Wild-type strain, isolated from petunia in NY, 2021 | This work |
| *Pectobacterium brasiliense* 1692 | Wild-type strain, isolated from potato in Brazil | (Duarte et al., 2004) |
| *Pectobacterium carotovorum* subsp. *carotovorum* WPP14 | Wild-type strain, isolated from potato in WI, 2001 | (Yap et al., 2004) |
| *Pectobacterium parmentieri* NY1722A | Wild-type strain, isolated from potato in NY, 2017 | (Ma et al., 2021) |
| *Pectobacterium versatile* NY1715C | Wild-type strain, isolated from potato in NY, 2017 | (Ma et al., 2021) |
| APA752 | Barcoded *mariner* vector pKMW3 ($Kan^R$) in *E. coli* WM3064 | (Wetmore et al., 2015) |
| APA766 | Barcoded *Tn5* vector pKMW3 ($Kan^R$) in *E. coli* WM3064 | (Wetmore et al., 2015) |
| Par6119 | *Pectobacterium aroidearum* 61-19 Tn5 transposon mutant library ($Kan^R$) | This work |
| PatSCRI1043 | *Pectobacterium atrosepticum* SCRI1043 *mariner* transposon mutant library ($Kan^R$) | This work |
| Pbr123 | *Pectobacterium brasiliense* 123-1 *mariner* transposon mutant library ($Kan^R$) | This work |
| Pbr1692 | *Pectobacterium brasiliense* 1692 *mariner* transposon mutant library ($Kan^R$) | This work |
| PccWPP14 | *Pectobacterium carotovorum* subsp. *carotovorum* WPP14 *mariner* transposon mutant library ($Kan^R$) | This work |
| PpaNY1722A | *Pectobacterium parmentieri* NY1722A Tn5 transposon mutant library ($Kan^R$) | This work |

| | | |
|---|---|---|
| PveNY1715C | *Pectobacterium versatile* NY1715C *mariner* transposon mutant library (Kan$^R$) | This work |

Table citations:
(Hinton et al., 1989; Duarte et al., 2004; Yap et al., 2004; Wetmore et al., 2015; Ma et al., 2021)

Supplementary Table 7. Statistics of the Nanopore read sets used to assemble genomes of the four strains novel to this study. The "Theoretical Coverage" is the ratio of the total number of raw bases to the length of the assembled genome. The "Actual Coverage" is the number of raw bases that actually align to the assembled genome. This is computed by aligning the filtered reads to the finished genome using Minimap2 (Li, 2016) and counting the number of aligned bases using Samtools (Li et al., 2009).

| Strain | Num raw reads | Num raw bases | Min length | Max length | Median length | Mean length | N50 | L50 | Theoretical Coverage | Actual Coverage |
|---|---|---|---|---|---|---|---|---|---|---|
| *P. aroidearum* 61-19 | 422,530 | 4,317,361,116 | 100 | 240,027 | 5,409 | 10,217.88 | 20,539 | 57,533 | 879.3 | 810.9 |
| *P. brasiliense* 123-1 | 278,576 | 3,029,762,188 | 101 | 383,643 | 7,010 | 10,875.89 | 18,985 | 48,839 | 632.3 | 578.4 |
| *P. parmentieri* NY1722A | 344,958 | 1,652,118,276 | 500 | 76,437 | 3,111 | 4,789.33 | 7,708 | 62,535 | 326.9 | 308.5 |
| *P. versatile* NY1715C | 189,581 | 697,145,140 | 1 | 77,530 | 2,372 | 3,677.29 | 5,989 | 33,043 | 132.2 | 124.3 |

Supplementary Table 8. Accessions for whole-genome assembly read and genome deposits in NCBI's SRA and Genbank repositories. All deposits were under the BioProject PRJNA1471679.

| Strain | BioSample | SRA (Illumina) | SRA (Nanopore) | Genbank |
|---|---|---|---|---|
| *P. aroidearum* 61-19 | SAMN60462425 | SRR39751989 | SRR39751990 | JBZUGZ000000000 |
| *P. brasiliense* 123-1 | SAMN60462424 | SRR39751987 | SRR39751988 | JBZUHA000000000 |
| *P. parmentieri* NY1722A | SAMN60462427 | SRR39751985 | SRR39751986 | JBZUGX000000000 |
| *P. versatile* NY1715C | SAMN60462426 | SRR39751983 | SRR39751984 | JBZUGY000000000 |

Summary Table 9. NCBI SRA accessions for barcoded transposon insertion-site sequencing (RB-TnSeq) for mutant library mapping. All deposits were under the BioProject PRJNA1471679.

Summary Table 10. NCBI SRA accessions for barcode sequencing (BarSeq) reads used for fitness assays. All deposits were under the BioProject PRJNA1471679.

Supplementary Figure 1. Transposon insertion distributions as measured in the mutant library pools. Strain PpaNY1722A contains a 8.3 kb plasmid included as a separate scaffold here. Plots were generated using "ggplot2" (Wickham, 2016) in R.

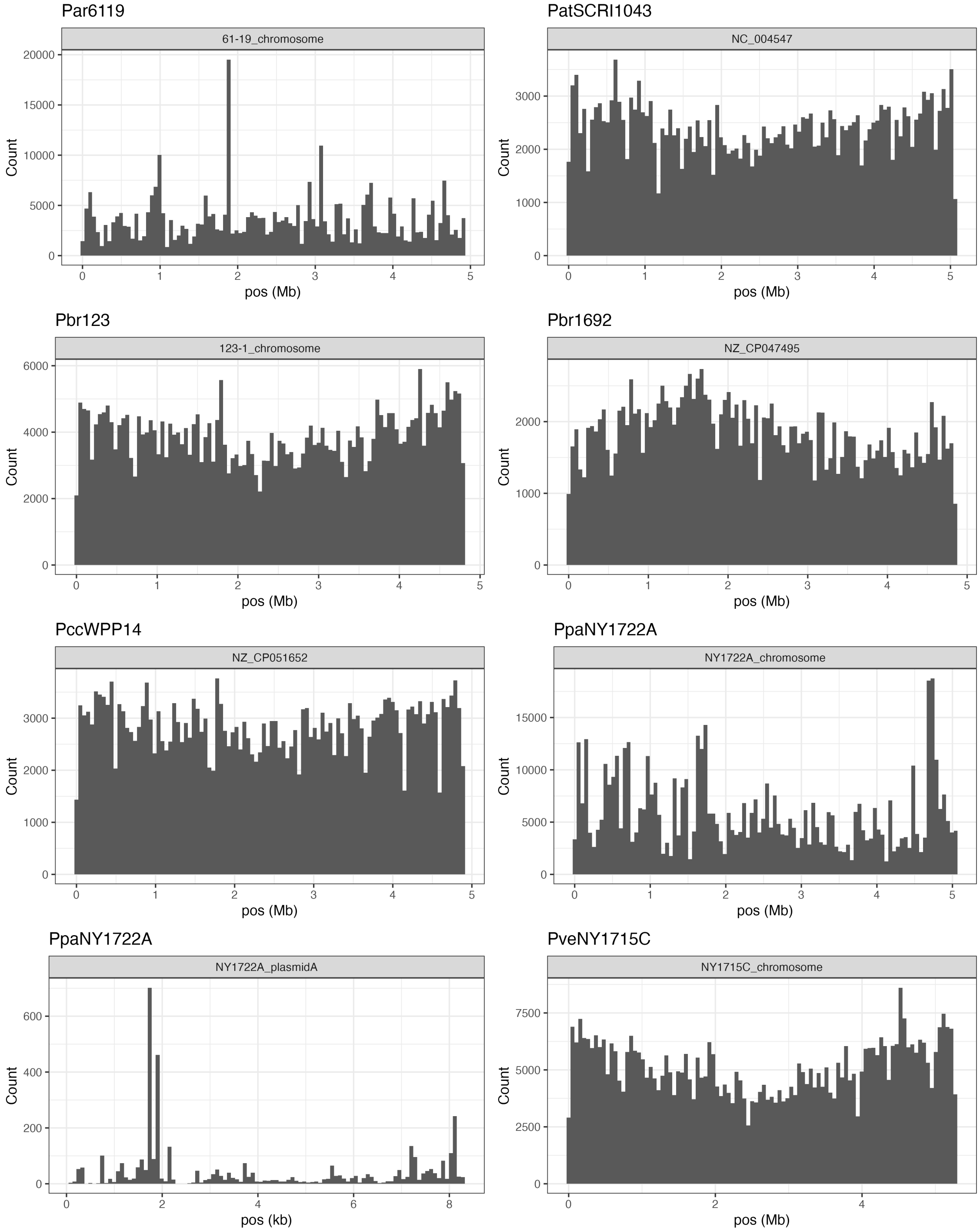

Supplementary Figure 2. UpSet plot of all (A) and predicted essential (B) orthologous genes across the seven strains tested here. UpSet plots were generated using “UpSetR” (Conway et al., 2017) in R.

A.

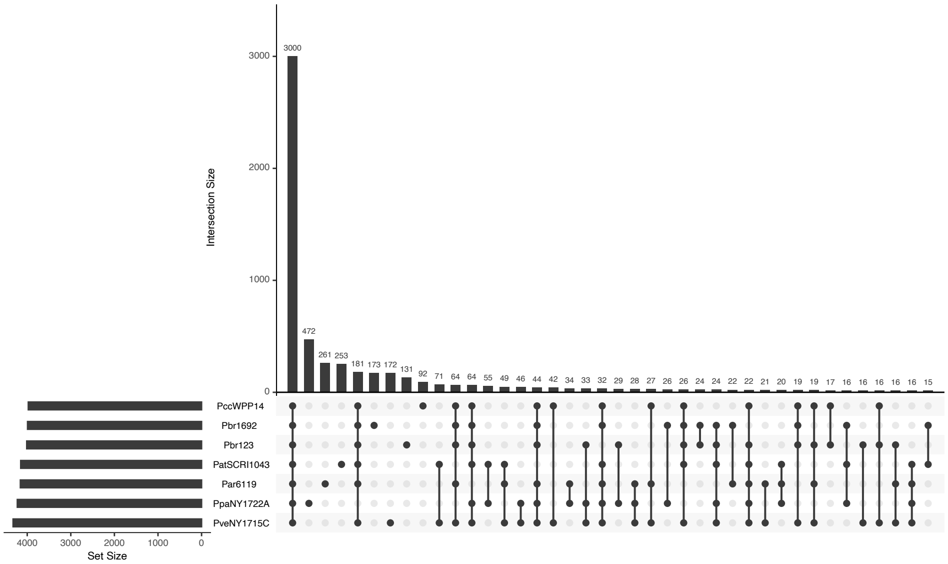


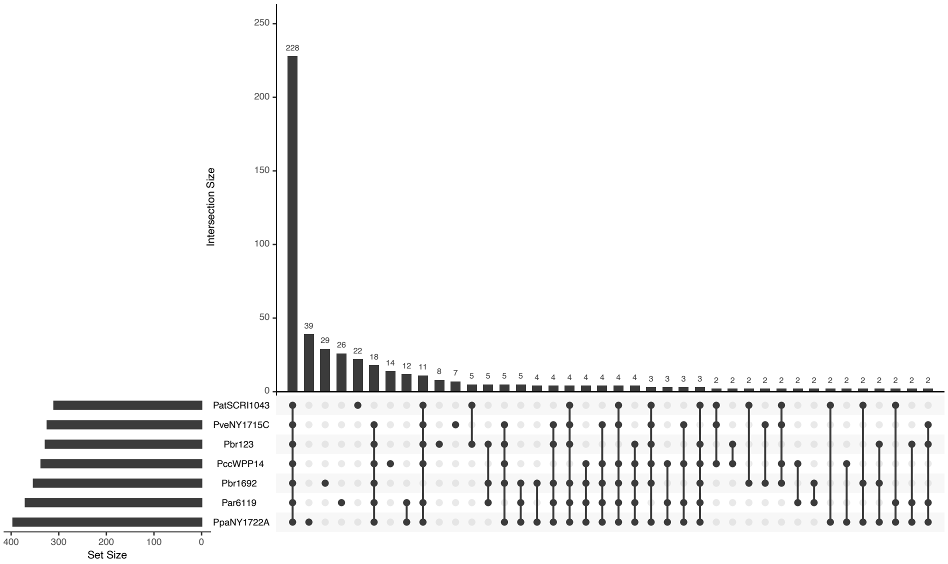


B.

Supplementary Figure 3. Principle component analysis of gene fitness scores for all strains, for experimental conditions tested that passed previously-established QC metrics. Plots were generated using “ggplot2” (Wickham, 2016) in R.

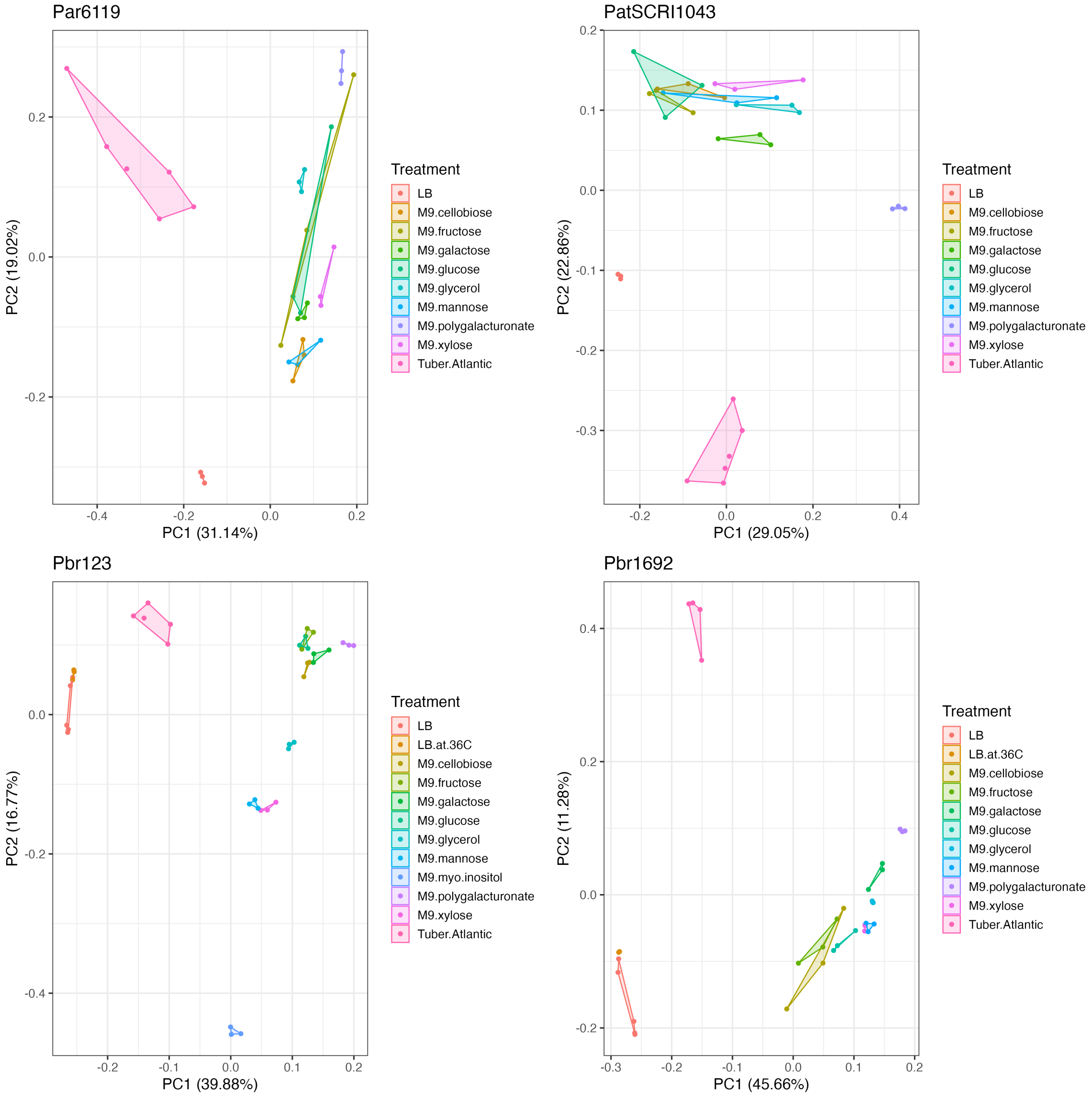

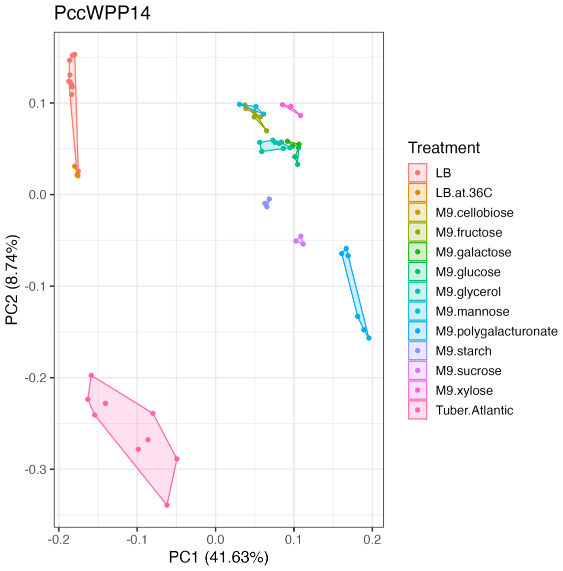
PccWPP14
PC2 (8.74%)
0.1
0.0
-0.1
-0.2
-0.3
-0.2
-0.1
0.0
0.1
0.2
PC1 (41.63%)
Treatment
LB
LB.at.36C
M9.cellobiose
M9.fructose
M9.galactose
M9.glucose
M9.glycerol
M9.mannose
M9.polygalacturonate
M9.starch
M9.sucrose
M9.xylose
Tuber.Atlantic

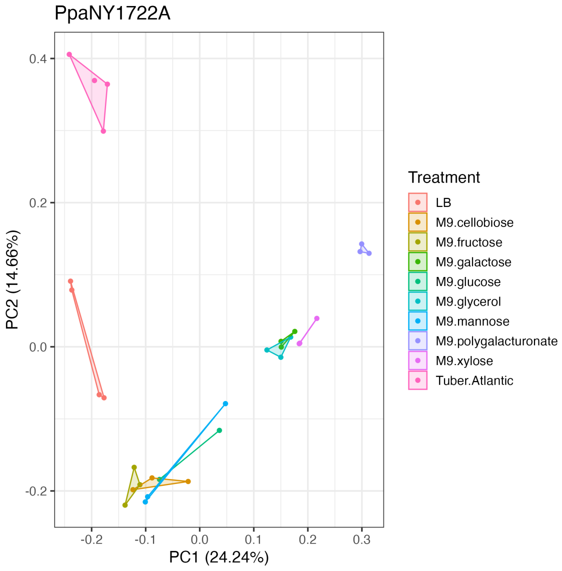
PpaNY1722A
PC2 (14.66%)
0.4
0.2
0.0
-0.2
-0.2
-0.1
0.0
0.1
0.2
0.3
PC1 (24.24%)
Treatment
LB
M9.cellobiose
M9.fructose
M9.galactose
M9.glucose
M9.glycerol
M9.mannose
M9.polygalacturonate
M9.xylose
Tuber.Atlantic

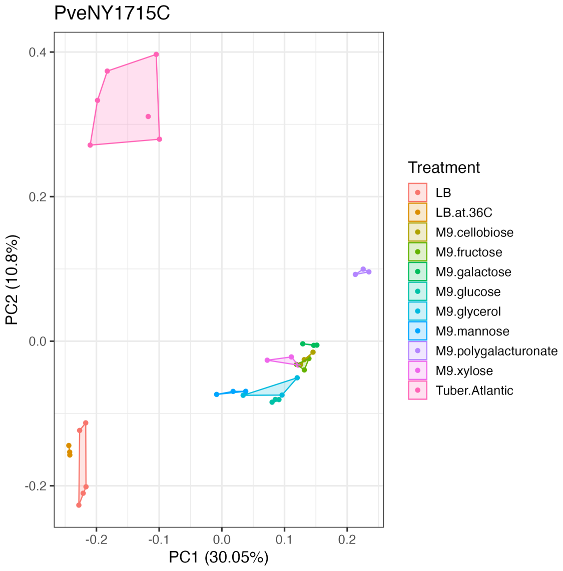
PveNY1715C
PC2 (10.8%)
0.4
0.2
0.0
-0.2
-0.2
-0.1
0.0
0.1
0.2
PC1 (30.05%)
Treatment
LB
LB.at.36C
M9.cellobiose
M9.fructose
M9.galactose
M9.glucose
M9.glycerol
M9.mannose
M9.polygalacturonate
M9.xylose
Tuber.Atlantic

Supplementary Figure 4. Mutations in the Enterobacterial Common Antigen (ECA) gene cluster decrease fitness of *Pectobacterium* strains across media and plant conditions. Gene fitness values shown for the 11 genes in the ECA. Heatmap was generated using “ggplot2” (Wickham, 2016) in R.

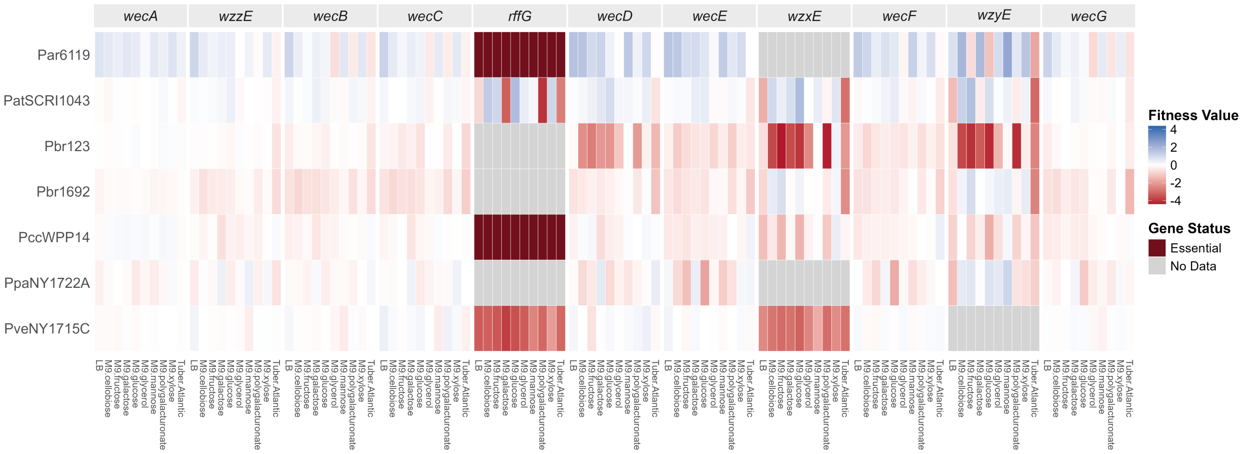

Supplementary Figure 5. Gene synteny visualized using clinker (Gilchrist and Chooi, 2021) of the group 4 capsule O-antigen cluster, from *wecA* (HER17_RS14615) to YjbH domain-containing protein encoding gene HER17_RS14485.

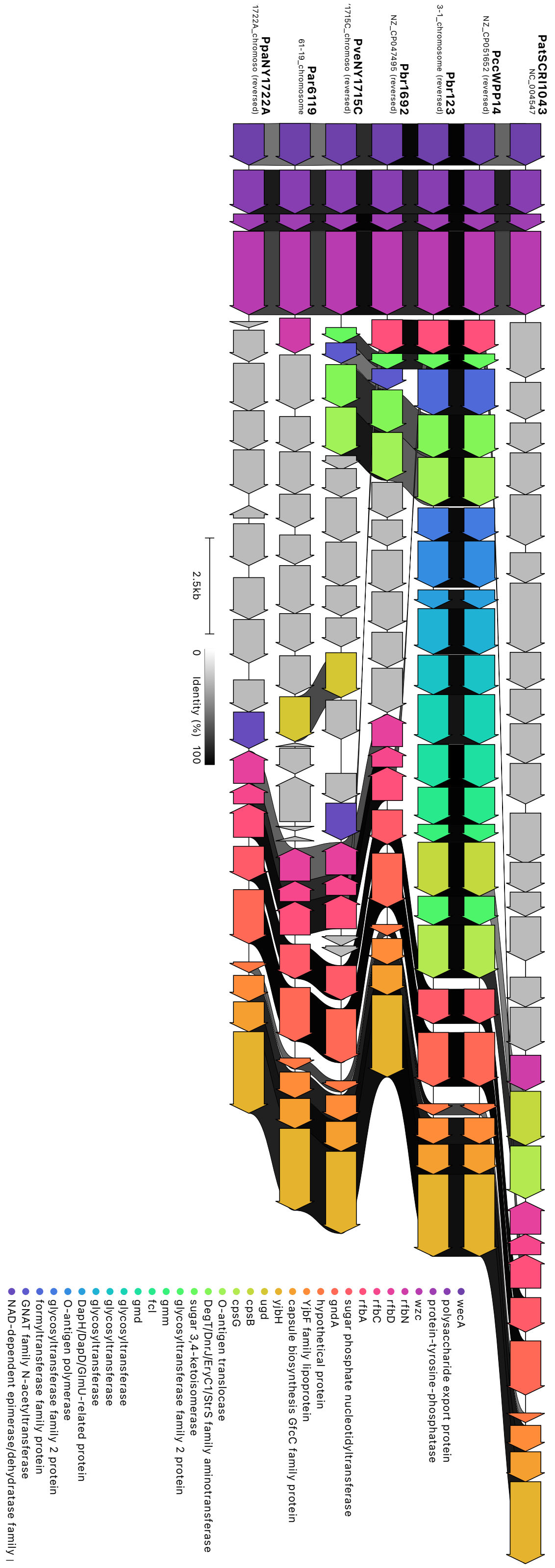
